# Pro-inflammatory feedback loops define immune responses to pathogenic lentivirus infection

**DOI:** 10.1101/2023.03.19.533358

**Authors:** Aaron J. Wilk, Joshua O. Marceau, Samuel W. Kazer, Ira Fleming, Vincent Miao, Jennyfer Galvez-Reyes, Alex K. Shalek, Susan Holmes, Julie Overbaugh, Catherine A. Blish

## Abstract

HIV causes chronic inflammation and AIDS in humans, though the rate of disease progression varies between individuals. Similarly, simian lentiviruses vary in their pathogenicity based on characteristics of both the host (simian species) and virus strain. Here, we profile immune responses in pig-tailed macaques infected with variants of SIV that differ in virulence to understand the immune mechanisms underlying lentiviral pathogenicity. Compared to a minimally pathogenic lentiviral variant, infection with a highly pathogenic variant results in a more delayed, broad, and sustained activation of inflammatory pathways, including an extensive global interferon signature. Conversely, individual cells infected with highly pathogenic lentivirus upregulated fewer interferon-stimulated genes at a lower magnitude, indicating that highly pathogenic lentivirus has evolved to partially escape from interferon responses. Further, we identified distinct gene co-expression patterns and cell-cell communication pathways that implicate *CXCL10* and *CXCL16* as important molecular drivers of inflammatory pathways specifically in response to highly pathogenic lentivirus infection. Immune responses to highly pathogenic lentivirus infection are characterized by amplifying regulatory circuits of pro-inflammatory cytokines with dense longitudinal connectivity. Our work presents a model of lentiviral pathogenicity where failures in early viral control mechanisms lead to delayed, sustained, and amplifying pro-inflammatory circuits, which has implications for other viral infections with highly variable disease courses.

## INTRODUCTION

Since the beginning of the HIV/AIDS pandemic, HIV has caused over 36 million deaths worldwide and remains a significant cause of morbidity and mortality globally (www.who.int). Chronic inflammation and CD4^+^ T cell loss are central to the pathogenesis of HIV^1–5^. In contrast, HIV’s simian counterparts, collectively referred to as simian immunodeficiency viruses (SIV), are highly variable in pathogenicity depending on both host genetic and viral strain factors, with many SIV strains being non-pathogenic in their natural hosts^3–5^. One of the most longstanding questions in lentiviral biology concerns the mechanistic basis for these starkly variable outcomes.

Virus-specific factors may play a role in differential lentiviral pathogenicity. For example, in simian precursors of HIV-1, the viral protein Nef has decreased activity in downregulating expression of CD3^6^. By promoting a greater degree of T cell activation upon infection, this modulation in Nef activity is thought to be a mechanism by which HIV-1 can cause chronic immune activation^7,8^. Additionally, simian precursors of HIV-1 also acquired the gene encoding Vpu, which enables escape from innate immune responses via evasion of tetherin and suppression of NF-κB signaling^9,10^.

The most direct evidence for virus-specific drivers of differential lentiviral pathogenicity comes from the study of emerging pathogenic variants of SIV where specific highly related variants were tested and shown to drive different clinical outcomes. It has long been known that genetic variants arise over the course of lentivirus infection that are antigenically and phenotypically distinct from the founder viruses present early in infection ^11^. For example, in pig-tailed macaques (*M. nemestrina*), genetic variants that evolve in the late stages of infection are highly cytopathic and rapidly replicating, unlike their precursors. By infecting immunocompetent pigtailed macaques with a SIV clone that has properties similar to transmitted forms of HIV, SIVMneCL8 (hereafter CL8) or a late-stage genetic variant that evolved from CL8 (SIVMne170, hereafter 170), Kimata, et al. showed that 170 was significantly more highly pathogenic than CL8 in pig-tailed macaques, thereby demonstrating that late-stage genetic variants were capable of directly driving disease progression(^12^.

These SIV variants that are closely related but show variable virulence thus represent a useful model to explore differential lentiviral pathogenicity within a host to define host virus-host interactions that drive pathogenesis. While the role of viral genetic determinants as the proximal cause of this differential pathogenicity have been described, the distal immune mechanisms resulting in these distinct outcomes remain undefined. Here, we performed longitudinal singlecell transcriptomics on peripheral blood mononuclear cells (PBMCs) from macaques infected either with CL8 or 170. We reveal broad and sustained pro-inflammatory responses as a defining feature of pathogenic lentiviral infection with the highly virulent variant, nominate potential molecular drivers of these pathways, and describe longitudinal cell-cell communication (CCC) networks distinguishing responses between non-pathogenic vs. pathogenic lentiviral infection. Collectively, our work provides a model for understanding beneficial vs. deleterious immune responses to lentivirus infection.

## RESULTS

### A transcriptomic atlas of immune responses to lentiviral pathogenic variants

To investigate the immune mechanisms that distinguish animals infected with variants of SIV that differ in their pathogenic properties, we performed Seq-Well-based(^13,14^ massively-parallel single-cell transcriptomic profiling on longitudinally-sampled PBMCs from pig-tailed macaques infected with either CL8 (*n* = 2; shades of blue) or 170 (*n* = 2; shades of red). As it is known that features of initial pathology in the days and weeks immediately after infection can predict overall disease outcome and tissue damage^15,16^, we profiled PBMCs from pre-infection and 8 postinfection time points, focusing on the hyperacute and acute stages of infection (**Figure 1A**). CL8 is a non-syncytium inducing SIV with slow replication kinetics; 170, conversely, is derived from an animal with late-stage infection with CL8, and is rapidly-replicating and syncytium inducing (**Figure 1B**). Compared to CL8-infected animals, animals infected with 170 have a peak viremia that is 6-fold higher and one week delayed and have a viral set point that is 3 orders of magnitude higher in subacute and chronic infection (**Figure 1C**)^12^. One animal, 170 B, died from simian AIDS-related complications at 31 weeks post-infection.

**Figure 1:**
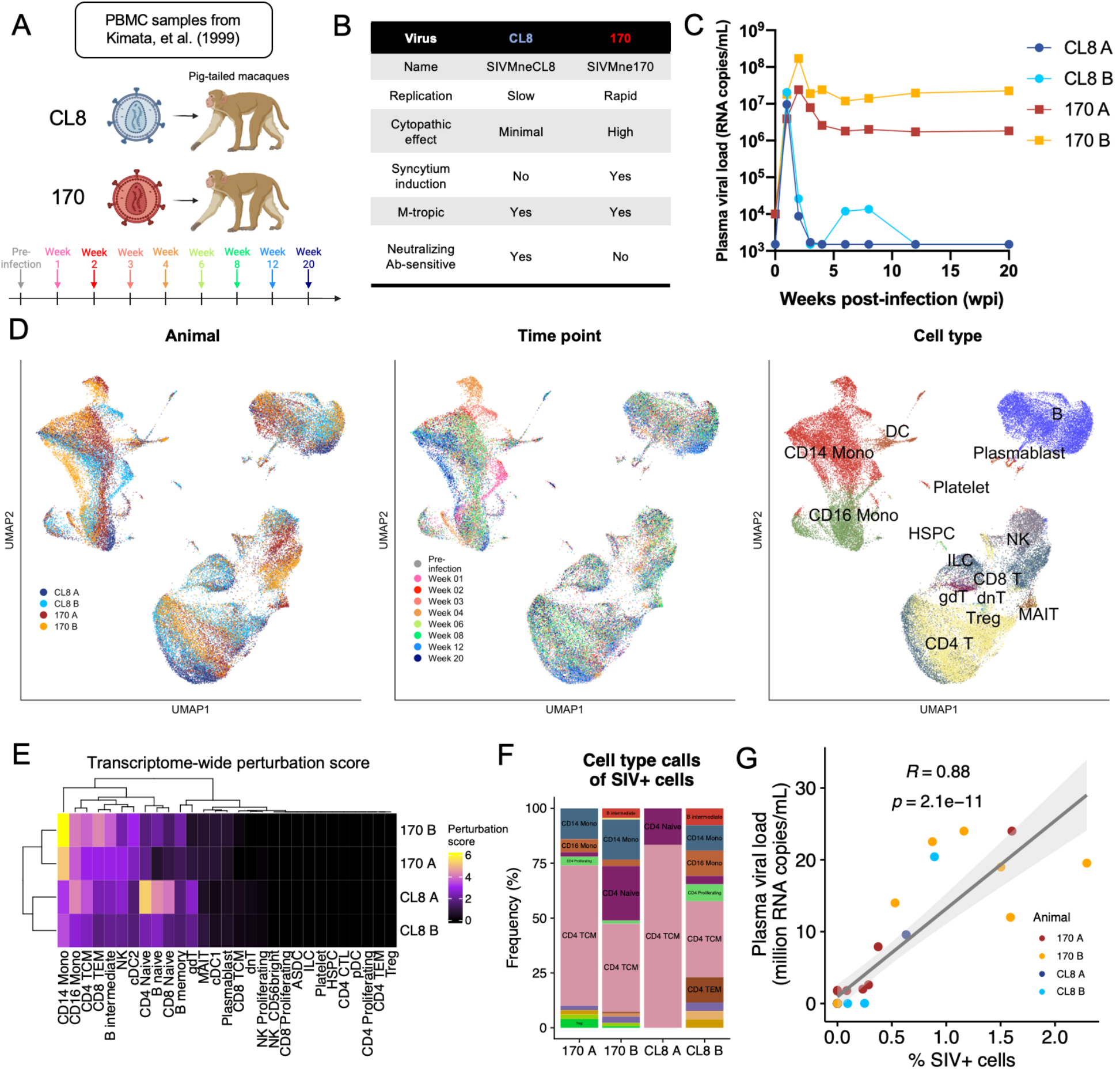
A single-cell atlas of immune responses to SIV pathogenic variants. **A)** Schematic of sample collection strategy. Pig-tailed macaques were infected with CL8 (shades of blue) or 170 (red/orange) and PBMCs sampled at the indicated time points. **B)** Virological and pathological characteristics of SIV pathogenic variants. **C)** Plasma viral loads of CL8- and 170-infected animals at the profiled time points. Animal 170 B died at 31 weeks post-infection from simian AIDS-related complications. **D)** UMAP projections of full scRNA-seq dataset colored by animal (left), time point (middle), and coarse cell type annotation (right). **E)** Heatmap of whole transcriptome perturbation score per cell type per animal, as described by^17^. **F)** Bar plot depicting cell type proportions of cells with SIV-aligned reads. **G)** Scatter plot depicting association between plasma viral load and the percentage of SIV^+^ cells in the scRNA-seq dataset.

In total, we sequenced 56,554 cells with an average of 1,570 cells per animal per time point (**Supplementary Figure 1**), which we aligned to a *M. nemestrina* reference genome that contained the viral genome sequences of CL8 and 170 in order to detect cells infected with SIV. Samples from 170-infected animals at 1 week post-infection had low viability and few high-quality cells were sequenced from these samples (**Supplementary Figure 1**; see **Methods**). Next, we created a merged feature matrix of all profiled samples that we subjected to dimensionality reduction by uniform manifold approximation and projection (UMAP), graphbased clustering, and cell type annotation (see **Methods**). UMAP projections of the full dataset indicated regions of the gene expression manifold that were strongly separated between time points and between 170- and CL8-infected animals, particularly within the myeloid compartment (**Figure 1D**). For example, we note a transcriptionally-unique population of monocytes that is composed almost entirely of cells from both 170-infected animals at 4 weeks post-infection, as well as distinct populations of monocytes from both CL8-infected animals at 1 week postinfection (**Figure 1D**, **Supplementary Figure 2**; see **Figure 3**). It is unlikely that this unique population of monocytes is related to a batch effect because other cell subsets from that time point, including T and B cells, appear well integrated in dimensionality reduction embeddings.

**Figure 2:**
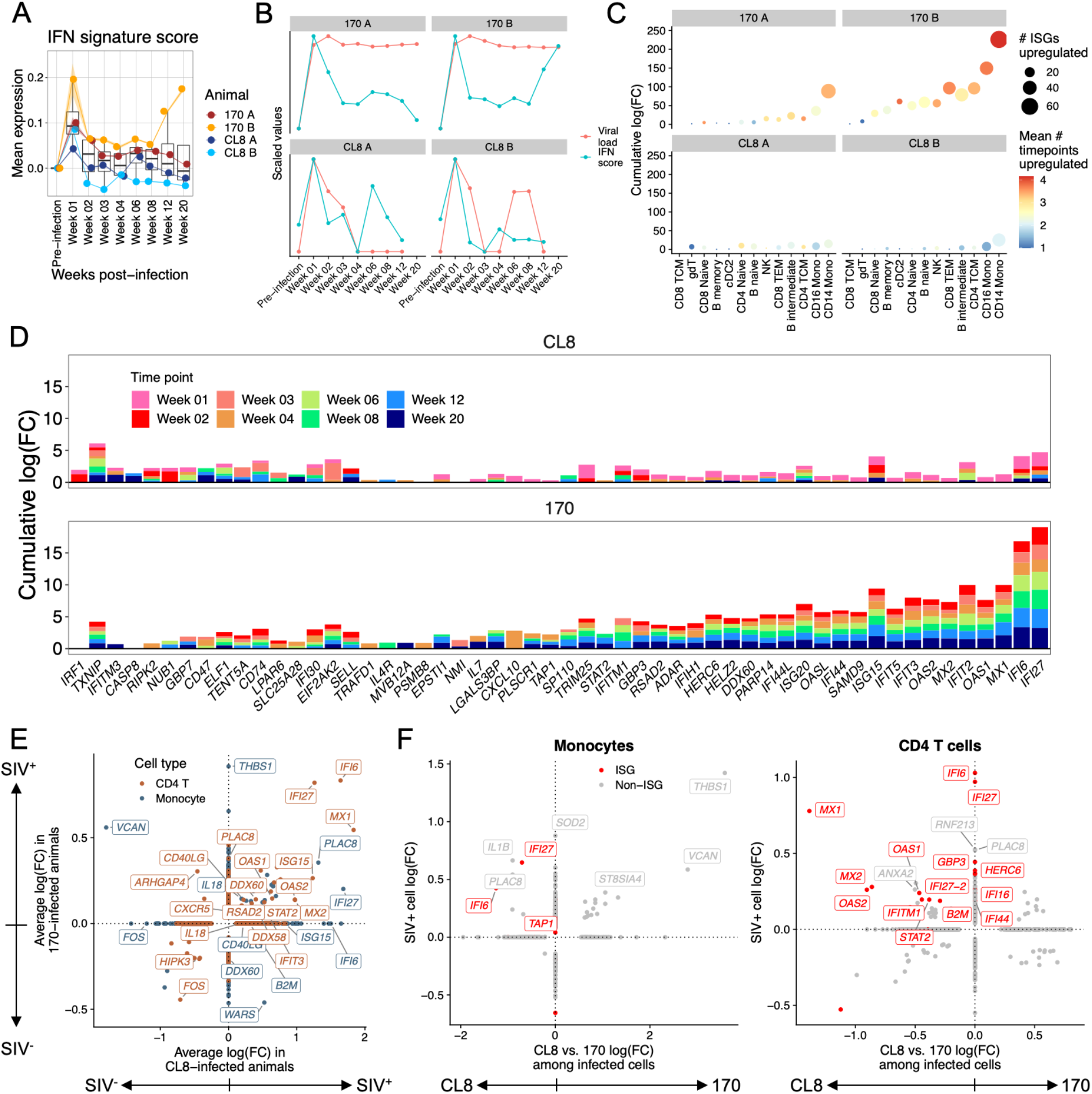
Broad and sustained interferon signatures in pathogenic SIV infection. **A)** Boxplot depicting mean expression of all ISGs over time across all profiled cells. **B)** Correlation between global IFN signature and viral load in each animal. **C-D)** For each animal, within each cell type, and at each non-baseline time point, differentially-expressed ISGs were calculated relative to pre-infection cells. **C)** For each ISG, we calculated the mean number of time points at which that ISG was significantly upregulated (color of points). For each cell type we calculated the number of ISGs significantly upregulated at any time point (size of points), and the cumulative log(fold-change) in expression of all significantly upregulated ISGs (*y*-axis). **D)** For each ISG, animal class (170 or CL8), and cell type, we calculated the cumulative log(fold-change) of significantly upregulated ISGs across all cell types. Only ISGs with a cumulative log(fold-change) of >0.75 in at least one animal group are shown. **E)** In the two SIV-infected cell types (monocytes and CD4^+^ T cells), we calculated DEGs between SIV^+^ cells relative to SIV^-^ cells in CL8 animals (*x*-axis) and 170 animals (*y*-axis). **F)** In monocytes (left) and CD4^+^ T cells (right), we calculated DEGs between SIV^+^ cells relative to SIV^-^ cells (*y-*axis). Then, among the SIV^+^ cells we calculated DEGs between cells from 170-infected animals and cells from CL8-infected animals (*x*-axis). For (**C-F**), all genes shown are significantly differentially expressed along at least one axis by Seurat’s implementation of the Wilcoxon rank-sum test.

**Figure 3:**
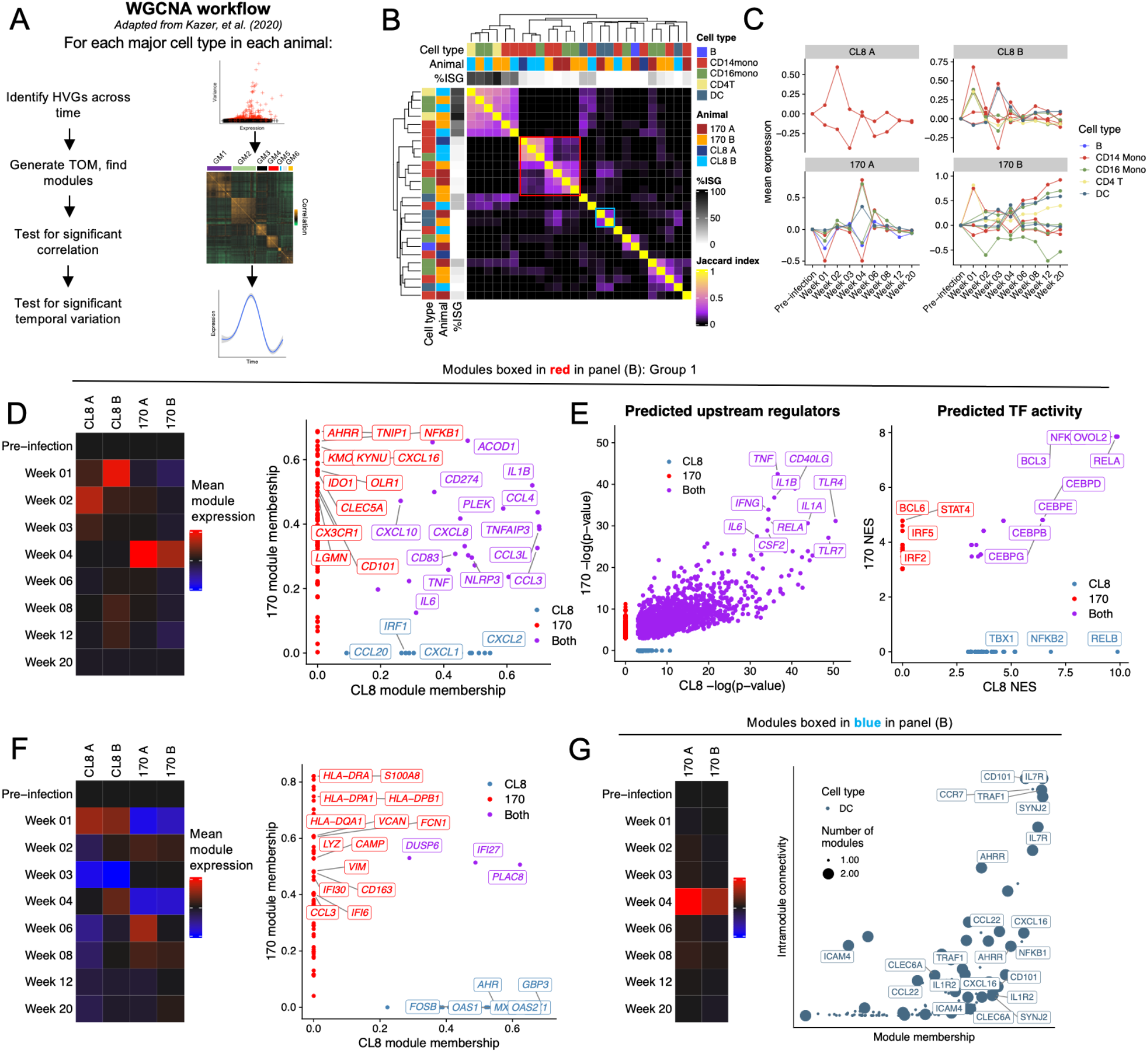
Gene module discovery reveals shared and distinct temporal features between immune responses to pathogenic and non-pathogenic lentiviral infection. **A)** Overview of gene module discovery by WGCNA, adapted from Kazer, et al.^22^. **B)** Jaccard index heatmap depicting the degree of overlap in gene membership between all discovered modules. **C)** Expression of modules over time by the cell type and in the animal in which the modules were discovered. For (**D-G**), heatmaps depict average expression of gene modules by the cell type and animal in which the modules were discovered. Scatter plots depict either the module membership of module genes in CL8 and 170 animals (right **D**, **F**, and **G**), or upstream regulators (left **E**) and transcription factor activities (right **E**) predicted to result in expression of the observed gene modules. **D-E)** monocyte modules boxed in red in panel (**B**). **F)** modules downregulated by monocytes in at least one time point. **G)** DC modules boxed in blue in panel (**B**).

To formally quantify the degree of this transcriptional perturbation, we calculated a genomewide perturbation score for each cell type relative to each animal’s pre-infection sample (see **Methods**). This perturbation score is calculated by first identifying genes that display evidence of differential expression between pre- and post-infection time points, calculating the difference of pseudobulk expression vectors of these genes between pre- and post-infection time points, and finally projecting the whole transcriptome of each animal onto this vector. This score thereby represents the magnitude of longitudinal whole-transcriptome shifts in gene expression, and reveals a strong longitudinal perturbation of CD14 monocytes in both 170-infected animals, as well as perturbation of CD16 monocytes and CD8^+^ T cell subsets (**Figure 1E**).

We next examined our ability to detect peripheral immune cells that were infected with SIV. The proportion of SIV^+^ cells ranged from 0-2.5% per sample and were predominantly found in memory CD4 T cells and monocytes, matching biological expectations (**Figure 1F**). SIV^+^ cells in CL8-infected animals were detected mainly at weeks 1-2 post-infection, while 170-infected cells were also present at later time points (**Supplementary Figure 1**). The distribution of SIV unique molecular identifiers (UMIs) in each infected cell was similar for CL8- and 170-infected cells (**Supplementary Figure 1**). The proportion of SIV^+^ cells correlated strongly with plasma viral load measurements (Pearson’s *r* = 0.88), suggesting the accurate identification of SIV^+^ cells (**Figure 1G**; discussed further in **Figure 2**).

To provide complementary validation to our single-cell transcriptional data, we also performed bulk RNA sequencing of all animals from pre-infection through 4 weeks post-infection (**Supplementary Figure 3**). This analysis revealed strong transcriptional perturbation in 170-infected animals that was most pronounced at 4 weeks post-infection, whereas transcriptional perturbation was largely resolved by 4 weeks post-infection in CL8-infected animals (**Supplementary Figure 3**). Additionally, these transcriptional responses were largely congruent between animals infected with the same virus. While we had very poor recovery of viable cells at 1 week post-infection in 170-infected animals, we were able to generate high-quality bulk transcriptomic data from one 170-infected animal at 1 week post-infection. Compared to pre-infection, the 1 week post-infection sample showed strong upregulation of interferon (IFN)-stimulated genes (ISGs; including *GBP3, MX2, DDX58*, and *APOBEC3B*) and pro-inflammatory chemokines *CCL8* and *CXCL10*, as well as platelet markers *PPBP, ALAS2, CAVIN2*, and *PRR35*. This sample also showed downregulation of cell type-specific marker genes including *CCR7, CD8B, GZMK*, and *CD14*, suggestive of a composition shift with decreased monocytes and lymphocytes and increased platelets (**Supplementary Figure 3**).

### Differential responses to interferon in minimally pathogenic vs. highly pathogenic SIV infection

As ISGs act as important restriction factors in acute infection^18–20^, we first analyzed how the longitudinal IFN response differed between CL8- and 170-infected animals. We established an IFN signature score composed of known ISGs (see **Methods**) and scored all cells in the dataset by expression of this signature. This revealed a kinetically dynamic IFN response that peaked in all animals at 1 week post-infection but stayed persistently elevated in 170-infected animals relative to CL8-infected animals (**Figure 2A**). Additionally, in 170 B, the global IFN signature increased monotonically from 8 weeks post-infection preceding death at 31 weeks post-infection (**Figure 2A**). Generally, the IFN signature was most concordant with viral load in both CL8 animals and highly discordant in 170 B, where the late increase in IFN signature is not paralleled by an increase in viral load (**Figure 2B**). This may either be the reflection of robust infection in another tissue, such as the intestine, or be the result of non-lentiviral opportunistic infections or ongoing non-infectious inflammatory stimuli.

To examine potential sources of type I IFN, we analyzed the expression of the upstream regulator of IFN, *IRF7*, as type I IFN-encoding genes themselves frequently are not detectable at the RNA level^21,22^. We found that global expression of *IRF7* generally correlated well with the global IFN signature and that plasmacytoid dendritic cells (pDCs) were the most prolific source of *IRF7* expression, even when accounting for their relative scarcity (**Supplementary Figure 4**). Additionally, in animal 170 B we found that the late increase in IFN signature was accompanied by a shift of *IRF7* expression from pDCs to conventional DCs and monocytes (**Supplementary Figure 4**), implicating these cells in sustaining IFN responses preceding death in this animal.

Next, we analyzed the breadth and duration of the IFN response. Both 170-infected animals upregulated substantially more ISGs than both CL8-infected animals, and these ISGs were upregulated at greater magnitude and at more time points in 170-infected animals (**Figure 2C**). Monocytes were the cell type that upregulated the greatest magnitude of ISGs in 170-infected animals, but cell type distribution of ISG upregulation was more evenly distributed in CL8-infected animals (**Figure 2C**). We also observed stark differences in which ISGs were expressed by 170-vs. CL8-infected animals. In 170-infected animals, the most upregulated ISGs were *IFI27, IFI6, MX1*, and *ISG15*, while in CL8-infected animals, *TXNIP* and *IFI30* were the most upregulated (**Figure 2D**). Several ISGs, including *MX1, MX2*, and *OAS1*, were upregulated in 170-infected animals at many different time points, but were generally only significantly upregulated in CL8 animals at weeks 1 and 2 post-infection (**Figure 2D**). There were no cell type specific patterns in ISG upregulation (**Supplementary Figure 5**). The ISG response in animal 170 B diversified in the later time points profiled, with ISGs including *IFI44L, RSAD2*, and *DDX60* comprising a larger proportion of the IFN response relative to previous time points (**Supplementary Figure 5**).

As ISGs can act as restriction factors during infection, we next examined how gene expression profiles differed between infected vs. bystander cells in 170-vs. CL8-infected animals. In each animal, we calculated differentially expressed (DE) genes between SIV^-^ and SIV^+^ cells in the two cell types that were the targets of infection, CD4^+^ T cells and monocytes. In SIV^+^ cells from all animals, we found upregulation of several ISGs in both cell types, consistent with autocrine IFN signaling (**Figure 2E**). Generally, ISGs were more broadly upregulated in infected CD4^+^ T cells compared to monocytes. For example, CD4^+^ T cells, but not monocytes, upregulated *OAS1, OAS2*, and *DDX58* upon infection in all animals (**Figure 2E**). We also observed broader ISG upregulation in infected cells from CL8-infected animals: in infected monocytes, *IFI6* and *ISG15* were only upregulated in CL8-infected animals, and *DDX60* was significantly downregulated in infected monocytes from 170-infected animals (**Figure 2E**).

While this analysis indicates which genes are upregulated in infected vs. bystander cells, we next sought to analyze how the magnitude of this upregulation differs between 170-vs. CL8-infected cells. To address this question, we calculated DE genes between SIV^+^ cells from CL8-infected animals and SIV^+^ cells from 170-infected animals. In both monocytes and CD4^+^ T cells, CL8-infected cells upregulated ISGs to a greater extent than 170-infected cells (**Figure 2F**), suggesting deficient expression of ISGs upon infection with SIVMne170. Since 170-infected animals have more SIV-infected cells at later time points relative to CL8-infected animals, we also assessed if the DEG results between CL8- and 170-infected animals were driven by infection kinetics. However, we found that 170-infected cells from 4 weeks post-infection and later expressed slightly higher magnitudes of some ISGs relative to their hyperacute infection counterparts (**Supplementary Figure 5**). This indicates that the lower expression of ISGs in 170-infected cells vs. CL8-infected cells (**Figure 2F**) is not confounded by infection kinetics. Collectively, this analysis reveals that *global* IFN signatures are broader and more sustained in highly pathogenic lentiviral infection (**Figures 2A-D**), but that minimally pathogenic lentivirus induces broader and greater expression of ISGs *within infected cells* (**Figures 2E-F**).

### Gene module discovery reveals structured longitudinal immune responses to infection with lentiviral pathogenic variants

To understand how cell transcriptional phenotype varies across time, we analyzed coordinated shifts in gene expression within each cell type that varied in time. We adapted a weighted gene correlation network analysis (WGCNA)-based^23^ approach to identify temporally-variable modules of co-expressed genes that was originally described by Kazer, et al.^22^. For each major cell type within each animal, we identified variable genes and used the single-cell gene expression matrices to generate topological overlap matrices (TOM) that capture gene co-expression structure. Modules of co-expressed genes were identified in these TOMs, and the correlation structure of each module was tested for statistical significance. Single cells were then scored for expression of each module, and module expression tested for significant variability across time (**Figure 3A**; see **Methods**). We perform this analysis a) for each cell type individually, so that temporally-variant gene modules are more likely to represent shifts in cell state rather than changes in cell type proportion and, b) for each animal individually, in order to be robust to inter-individual heterogeneity. These gene modules thus represent coordinated programs of cellular phenotype and can be used to nominate cell type-specific drivers of synchronized responses to infection.

This process returned 26 distinct, temporally-variant gene modules in B cells, monocytes, CD4 T cells, and DCs (**Figure 3B**). No significantly-correlated, temporally-variant gene modules were discovered in CD8 T cells or NK cells, suggesting a lack of temporally coordinated shifts in gene program expression over time. To visualize relationships between the identified gene modules, we calculated the Jaccard index between each set of modules, which revealed groups of partially overlapping modules discovered between distinct animals or cell types (**Figure 3B**). For example, gene module discovery revealed a set of 6 highly overlapping modules that were composed primarily of ISGs (**Figure 3B**); expression of two of these modules were highly significantly enriched in SIV^+^ cells (**Supplementary Figure 6**). We next scored the cells of each animal and cell type for expression of the corresponding gene modules to examine how expression of each module varied across time. This revealed shifts in module expression that paralleled global patterns in whole-transcriptome perturbation (**Figure 1D-E**), where 170-infected animals display concordant shifts in monocyte phenotype at 4 weeks post-infection (**Figure 3C**).

Next, we analyzed a group of 7 partially overlapping modules that were discovered within monocytes of all four animals (**Figure 3B**; red box, hereafter “Group 1”). Expression of the corresponding modules peaked at 1-2 weeks post-infection in CL8-infected animals, but at 4 weeks post-infection in both 170-infected animals (**Figure 3D**). Genes that were common to all Group 1 modules included pro-inflammatory cytokines and chemokines *IL1B, TNF, CCL3, CCL4, IL6, CXCL8*, and *CXCL10*, as well as inflammasome component *NLRP3*. Group 1 modules from CL8-infected animals contained the neutrophil-chemoattractants *CXCL1* and *CXCL2*, while modules from 170-infected animals contained the chemokine *CXCL16* (**Figure 3D**). Additionally, modules from 170-infected animals contained three separate members of the kynurenine pathway of tryptophan metabolism (*IDO, KYNU*, and *KMO;* **Figure 3D**); activity of this pathway is known to inhibit T cell proliferation and IL-17 production and strongly predict HIV mortality^24,25^. We performed ingenuity pathway analysis (IPA) to identify predicted upstream regulators of these modules, as well as transcription factor (TF) activity analysis^26^ to identify potential regulatory mechanisms underlying module expression. These analyses implicated a similar set of upstream drivers in module expression, including *IL1B* and *IFNG*, but only partially overlapping TF activities, where TFs from the non-canonical pathway of NF-κB activation (NFKB2 and RELB) where predicted to drive expression of modules from CL8-infected animals but not 170-infected animals (**Figure 3E**). This paralleled higher expression of receptors that induce non-canonical NF-κB signaling *TNFRSF13C* (encoding BAFFR) and *TNFRSF11A* (encoding RANK) in CL8-infected animals (**Supplementary Figure 6**).

We noted there were monocyte-based modules that showed the opposite expression pattern and were predominantly repressed relative to baseline, often concomitant with upregulation of Group 1 modules (**Figure 3C, F**). In 170-infected animals, these predominantly repressed modules contain several HLA class II-encoding as well as the immunoregulatory hemoglobin scavenger *CD163*, and antimicrobial agent-encoding genes *LYZ* and *CAMP* (**Figure 3F**). Collectively, these results suggest a synchronized and coordinated shift from antigen presentation and antimicrobial function towards pro-inflammatory cytokine and chemokine production in monocytes during highly pathogenic lentivirus infection.

We also identified two overlapping modules discovered in both 170-infected animals in DCs. These modules were expressed in both 170-infected animals at predominantly 4 weeks postinfection and contained genes implicated in DC trafficking to secondary lymphoid tissues, including *CCR7* and *IL7R^27^*, pro-inflammatory chemokine *CXCL16*, and molecules involved in communication with T cells, including *CD101* and *CCL22^28,29^* (**Figure 1G**). As *CCR7* and *IL7R* are highly expressed on most T cells, we confirmed that *CCR7^+^IL7R^+^* DCs expressed canonical DC markers without co-expression of any canonical T cell-defining genes (**Supplementary Figure 6**). Collectively, gene module discovery analysis reveals coordinated pathways that distinguish immune responses between cell types, states, and lentiviral pathogenic variants.

### Single-cell resolution communication analysis reveals inflammatory networks differentiating immune responses between lentiviral pathogenic variants

While our gene module discovery analysis nominates potential drivers of shifts in cell state over time, we wished to formally analyze how communication pathways between cells, or cell-cell communication (CCC) influenced downstream transcriptional phenotypes. To address this question, we applied Scriabin, a method for comparative CCC analysis at single-cell resolution that does not require any degree of subsampling or aggregation (**Figure 4A**; see **Methods**)^30^.

**Figure 4:**
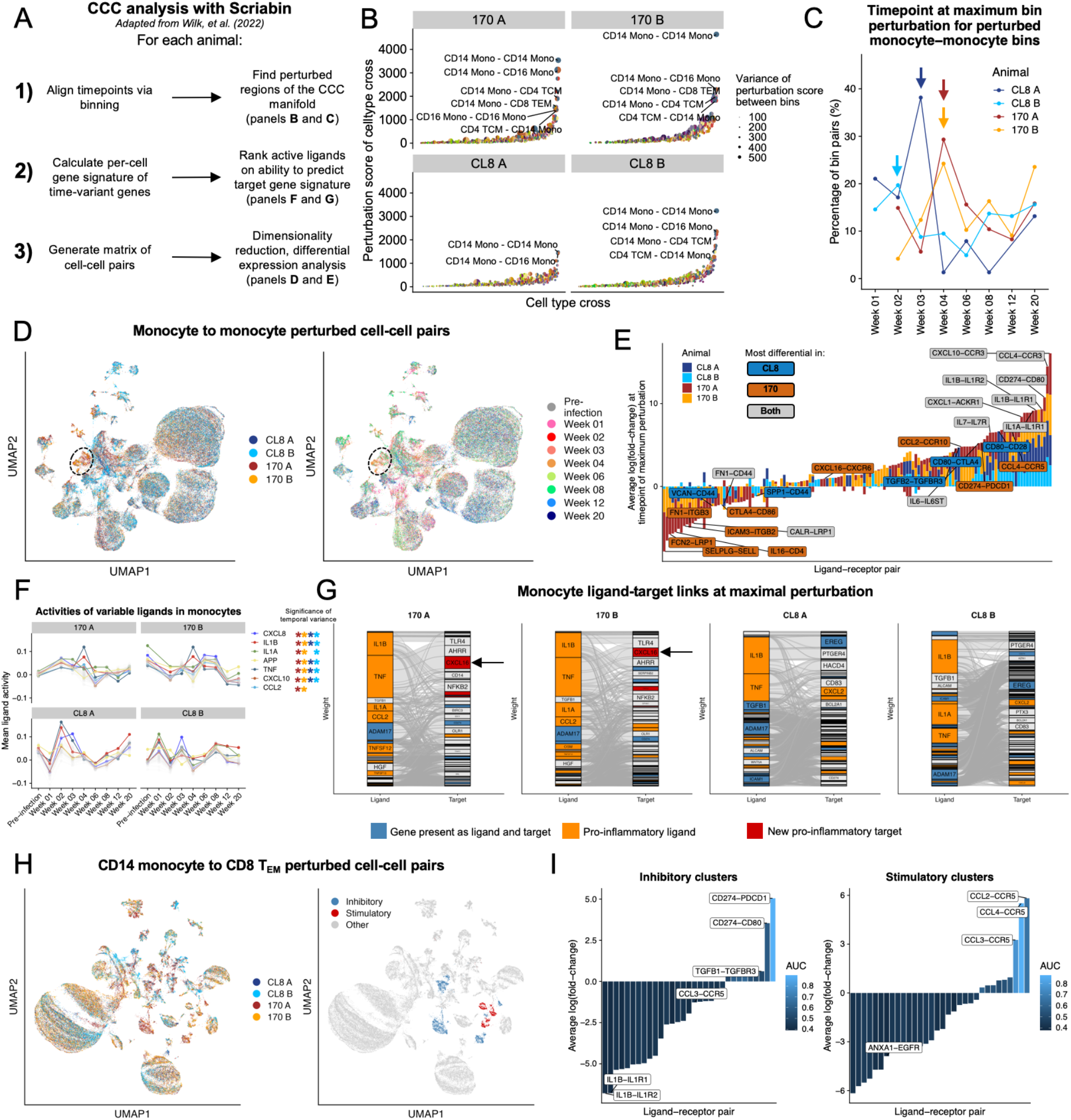
CCC pathways distinguishing immune responses to pathogenic and non-pathogenic lentiviral infection. **A)** Overview of application of Scriabin^30^ to analyze single-cell resolution CCC profiles. Briefly, Scriabin’s summarized interaction graph workflow was used to align the time points within an animal and identify cell type combinations with perturbed CCC to prioritize for downstream analysis. Then, Scriabin’s ligand activity ranking process was used to identify biologically active CCC edges. Finally, Scriabin’s CCIM workflow was used to visualize the full structure of CCC phenotypes (see **Methods**). **B)** Scatter plots depicting the degree of CCC perturbation predicted by Scriabin’s summarized interaction graph workflow (*y*-axis). The most highly perturbed cell type combinations for each animal are labeled (sender cell type – receiver cell type). Points are sized by the variability in perturbation score between bins of the same cell type combination. **C)** Plot depicting the distribution of time points at which CCC in each monocyte-to-monocyte bin-bin pair is maximally perturbed. **D)** UMAP projections of the most highly perturbed monocyte-monocyte cell-cell pairs from all samples, colored by animal (left) and time point (right). **E)** Stacked bar plot depicting DE ligand-receptor pairs in the monocytes of each animal at the time point of maximal perturbation. A ligand-receptor pair is considered to be more differentially expressed in an animal class if >1 standard deviation of that ligand-receptor pair’s log(fold-change) is contributed by a particular animal class. **F)** Predicted ligand activities of monocytes over time in each animal. **G)** Alluvial plots depicting ligand-target gene connections within monocytes of each animal at the time point of maximal perturbation. **H)** UMAP projections of the most highly perturbed CD14 monocyte-CD8 T_EM_ sender-receiver cellcell pairs colored by animal (left) and clusters enriched in 170-infected animals (right). **I)** Bar plots depicting DE ligand-receptor pairs in the clusters highlighted in (**H**).

To focus our CCC analysis on the cell types with the highest degree of CCC perturbation over time, we applied Scriabin’s summarized interaction graph workflow: for each sample, we generated a summarized interaction graph representing the total magnitude of potential interactions between every pair of cells. Then, we aligned datasets from all the time points from a given animal using Scriabin’s binning workflow, which assigns each cell a bin identity that maximizes the similarity of cells within each bin while simultaneously maximizing representation of all time points within each bin. These bin identities thus represent high-resolution intra-time point correspondences. We used our reference-based cell type annotations, which were transferred via neighbor graphs that are not used for binning, as orthogonal validation that binning recovered phenotypically similar cells (**Supplementary Figure 7**). We next assessed variability in total communicative potential across time within each bin, which is summarized in **Figure 4B**. This process revealed that monocyte-to-monocyte communication is the most longitudinally perturbed across all four animals, and additionally indicated that monocyte communication with memory T cells and NK cells were also significantly perturbed (**Figure 4B**). In 170-infected animals, the most highly perturbed monocyte-to-monocyte bins displayed the largest shift in CCC magnitude at 4 weeks post-infection, while CL8 animals had the greatest perturbation at 1-2 weeks post-infection (**Figure 4C**). For downstream analyses, we defined the time point of maximal perturbation as 4 weeks post-infection for 170-infected animals, 2 weeks post-infection for CL8 A, and 1 week post-infection for CL8 B.

To comprehensively visualize the CCC structure of our dataset, we next applied Scriabin’s cellcell interaction matrix (CCIM) workflow. This workflow generates a matrix of cell-cell pairs by ligand-receptor pairs that can be treated analogously to a gene expression matrix for downstream analytical tasks like dimensionality reduction and differential expression testing (**Figure 4A**). We also used Scriabin to predict which ligand-receptor pairs were biologically active by leveraging information about gene expression changes downstream of specific ligands, and incorporated this information into the CCIM (see **Methods**).

We first used Scriabin’s CCIM workflow to analyze the most highly perturbed CCC between cellcell pairs of monocytes. This analysis revealed groups of monocyte-monocyte pairs that appeared to have distinct communication profiles (**Figure 4D**). Differential expression testing indicated that CCC between monocytes at the time point of maximal perturbation was characterized by increased signaling through *IL1B* and *CCL4* (**Figure 4E**; **Supplementary Table 5**). 170-infected animals alone had increased signaling with *CXCL16* and *CCL2*, while losing communication through the adhesion molecule *SELPLG*. Conversely, monocytes from CL8-infected animals had increased communication via *TGFB2* (**Figure 4E**). These shifts in CCC pathways were also reflected in single-cell level ligand activities, which showed increased predicted activity of *IL1B* at the time point of maximal perturbation in all animals, but only an increase in *CCL2* activity at 4 weeks post-infection in both 170-infected animals (**Figure 4F**).

Next, we examined what downstream gene expression changes were predicted to result from the observed ligand activities. While *IL1B* and *TNF* were the primary predicted drivers of target gene profiles in all animals, in 170-infected animals these ligands were predicted to result in upregulation of *TLR4, NFKB2*, and the aryl hydrocarbon receptor repressor *AHRR*, but in CL8-infected animals were predicted to result in the upregulation of neutrophil chemoattractant *CXCL2* and checkpoint molecule *CD83* (**Figure 4G**)^31^. Intriguingly, in 170-infected animals alone we observed upregulation of additional pro-inflammatory cytokines and chemokines like *CXCL16* that had not been predicted to be active as ligands (**Figure 4G**), suggesting a broadening of pro-inflammatory signaling in infection with highly pathogenic lentivirus. Collectively, these results demonstrate overlapping yet distinct monocyte-to-monocyte communication mechanisms in response to lentiviral variants of different pathogenicity, with highly pathogenic lentivirus generally inducing a delayed and broader pro-inflammatory communication profile.

We next analyzed communication from CD14 monocytes to CD8 T_EM_ cells, as CCC for this sender-receiver cell type combination was highly perturbed only in 170-infected animals (**Figure 4B**). Using Scriabin’s CCIM workflow on highly perturbed CD14 monocyte-CD8 T_EM_ cell-cell pairs, we found several clusters that appeared highly enriched for cell-cell pairs from 170-infected animals (**Figure 4H**). We found that three of these clusters showed increased communication through inhibitory pathways, including the *CD274-PDCD1* (PDL1-PD1) pathway, while the other three clusters showed increased communication through chemokines *CCL2, CCL3*, and *CCL4* through their cognate receptor *CCR5* (**Figure 4I**). Interestingly, we found that the frequency of both groups of clusters were highest in the 170-infected animals at 4 weeks post-infection (**Supplementary Figure 8**). Thus, Scriabin revealed the concomitant use of opposing signaling modalities between CD14 monocytes and CD8 T_EM_ cells in response to highly pathogenic lentivirus infection, underscoring the utility of using single-cell resolution CCC methods to capture the full structure of CCC phenotypes.

### Broad longitudinal inflammatory networks in pathogenic lentiviral infection

While our single-cell resolution analysis of CCC pathways enabled us to compare what communication edges were differentially active between different animals and time points, we were next interested in formally analyzing how these communication pathways operate <u>across</u> time points in order to understand the dynamic evolution of CCC during infection. To address this question, we applied Scriabin’s longitudinal circuitry discovery workflow^30^. A longitudinal communicative circuit is a set of communication edges at two consecutive time points, where communication at the second time point is predicted to be the consequence of communication at the first time point (**Figure 5A**). Consider an interacting sender-receiver cell pair S_1_-R_1_ at time point 1 and interacting sender-receiver cell pair S_2_-R_2_ at time point 2. If communication between S1-R1 results in the expression of ligand L_A_ by R_1_, S_1_-R_1_-S_2_-R_2_ represents a communicative circuit if R_1_ and S_2_ occupy the same bin (i.e., S_2_ represents the phenotypic counterpart of R_1_ at time point 2) and if L_A_ is predicted to be an active ligand in the S_2_-R_2_ interaction (**Figure 5A**). Thus, longitudinal circuits represent the single-cell resolution stitching together of two neighboring time points by succeedent communication pathways.

**Figure 5:**
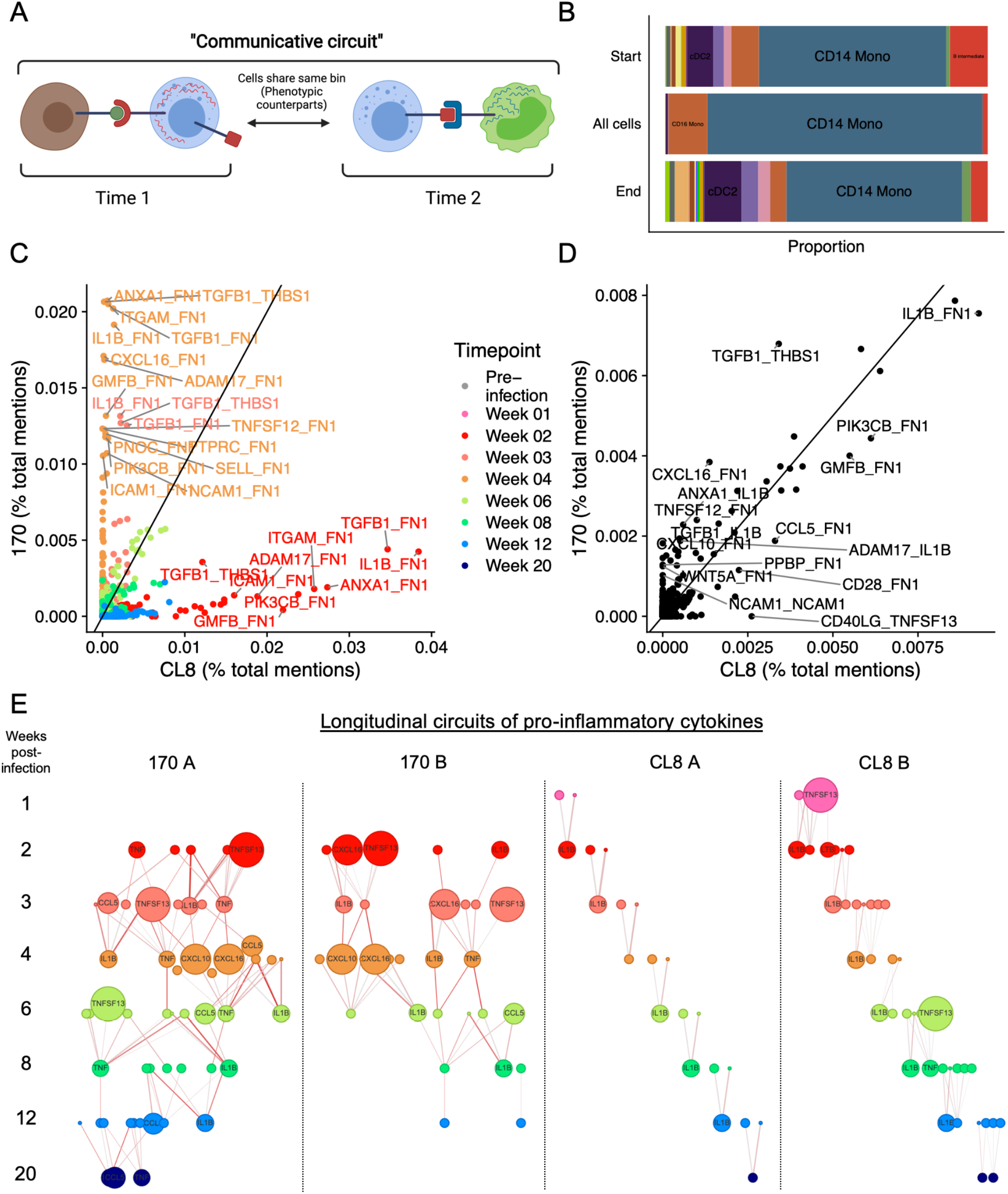
Longitudinal CCC circuits during immune response to lentiviral pathogenic variants. **A)** Diagram illustrating the definition of a longitudinal communicative circuit. **B)** Stacked bar plot depicting the proportion of cell type annotations for cells from animal 170 A participating in longitudinal circuits. Proportions are separated by cells participating at the beginning or the end of the circuit, as well as the total distribution of all cells participating in circuits. **C-D)** Scatter plots depicting ligand-target pairs participating in longitudinal communication circuits that differentiate 170-infected animals (*y*-axis) and CL8-infected animals (*x*-axis). For (**C**), each point represents a ligand-target pair participating in a circuit at a particular time point. For (**D**), each point represents the total number of mentions for each ligand-target pair in all circuits, regardless of time point. **E)** Longitudinal circuits of pro-inflammatory cytokines in each animal. Circuits were restricted to pro-inflammatory cytokines (see **Supplementary Table 4**) and networks of connected pro-inflammatory cytokine ligand-target pairs plotted for each animal across time. Points are sized by the expression of the respective cytokine at each time point. Edges are colored by the product of the ligand activity score and the target weight for the sequential cytokine (see **Methods**).

We applied Scriabin’s circuitry discovery workflow to identify all temporally consecutive communication circuits across time in each animal. We first noted that the vast majority of cells participating in circuits were monocytes; however, non-monocyte cell types were more highly represented in cells at the beginnings and ends of circuits (**Figure 5B**). This indicates that monocytes serve as the hubs and authorities of longitudinal communication networks, integrating and sending signals to other cell types to coordinate downstream responses.

We next examined which longitudinal circuits differed between 170-vs. CL8-infected animals. When analyzing ligand-target pairs participating in circuits at each time point, we observed strong differences between 170- and CL8-infected animals; most 170-infected animal circuits overlapped at the 4 week post-infection time point, compared to the 2 week post-infection time point for CL8-infected animals (**Figure 5C**). However, when summarizing the frequency of ligand-target pairs participating in circuits irrespective of time point, we found more concordant patterns between 170- and CL8-infected animals (**Figure 5D**). This indicates that longitudinal communication pathways in response to non-pathogenic vs. pathogenic lentiviral infection differ primarily by the time at which they operate rather than the specific ligands that compose these pathways. Nonetheless, we noted several ligand-target pairs that were more enriched in circuits from 170-infected animals, including *CXCL10* and *CXCL16* acting as ligands for the upregulation of extracellular matrix protein *FN1* (**Figure 5D**).

Our single-cell resolution analysis of CCC nominated putative signaling mechanisms for the expansion of the breadth of pro-inflammatory cytokine expression in 170-infected animals (see **Figure 4G**), and our analysis of longitudinal communication circuits identified two chemokines that were enriched in circuits from 170-infected animals. We hypothesized that this broader signature of pro-inflammatory cytokine activity in response to pathogenic lentivirus infection may reflect distinct network structures of cytokine regulation. To comprehensively visualize the longitudinal evolution of pro-inflammatory cytokine expression and regulation, we identified all communicative circuits involving sets of pro-inflammatory cytokines that were consecutive in time and regulation. This analysis revealed broad, sustained, and densely connected longitudinal networks of pro-inflammatory cytokine expression exclusively in 170-infected animals (**Figure 5E**). Networks from CL8-infected animals, conversely, were limited and sparsely connected across all time points. While circuits from both 170- and CL8-infected animals included *IL1B, TNF, TNFSF13*, and *CCL5*, only networks from 170-infected animals included the chemokine *CXCL16* (**Figure 5E**). Collectively, these results are suggestive of inflammatory feedback loops that are specific to immune responses to highly pathogenic lentivirus infection.

## DISCUSSION

In this work, we have applied single-cell transcriptomics and novel frameworks for CCC analysis to examine features of the immune response that distinguish infection with minimally vs. highly pathogenic lentivirus infection. A substantial body of work has compared differential lentiviral pathogenicity from the perspective of host genetics by infecting different non-human primate (NHP) species with SIVs from different natural hosts^4,32–36^. However, the role of virus-specific factors remains relatively underexplored. We leveraged unique samples that allowed us to compare responses at weekly intervals after infection to the gene profile at the time of exposure by leveraging a model of differential lentiviral pathogenicity where all hosts were from the same genetic background^12^, we here provide the first description of virus-specific immune profiles that distinguish minimally pathogenic vs. highly pathogenic lentivirus infection.

Compared to infection with minimally pathogenic CL8, we find that infection with highly pathogenic 170 induces delayed and broad signatures of immune activation. This finding is in line with evidence that chronic inflammation and immune activation is a distinguishing feature of pathogenic SIV infection as well as HIV infection relative to SIV infection in natural hosts^18,22,37,38^, but for the first time identifies densely connected longitudinal pro-inflammatory circuits originating from monocytes as a source of this inflammation. Prior work indicating that persistent type I IFN signaling in pathogenic SIV leads to broad dysfunction in both innate and adaptive immunity during chronic infection^37,39–43^. In line with these observations, here we show that 170-infected animals display a broad and sustained response to type I IFN and demonstrate for the first time that distinct sets of ISGs are upregulated in response to IFN in chronic and pathogenic infection. Similarly, preliminary evidence suggests blockade of type I IFN signaling during chronic infection may suppress other inflammatory pathways and pathology and rescue counts of HIV-specific T cells^39,42,44^. The association between delayed/sustained IFN signaling and disease outcome has also been postulated in the setting of acute viral infections like MERS-CoV and SARS-CoV-2^45–47^, indicating this may be a conserved feature of imbalanced or ineffective antiviral immunity.

Conversely, we found that infected CD4^+^ T cells and monocytes from animals infected with mildly pathogenic CL8 upregulated a broader range of ISGs and expressed a higher magnitude of these ISGs than the same cells in 170-infected animals. This finding is consistent with an evolved immune evasion mechanism of 170, where 170 may escape detection via decreased PRR sensitivity, IFN production, or autocrine IFN signaling. As ISGs can act as critical antiviral restriction factors in infected cells^19,22^, we hypothesize that this represents an evolved immune evasion strategy of 170 and may be an explanatory mechanism for the higher peak viremia and set point in 170-infected animals. Thus, while 170-infected animals have a broad *global* IFN signature, individual cells infected with 170 fail to upregulate the same set of ISGs to the same extent. These findings are specifically made possible by our single-cell approach and ability to accurately distinguish infected from bystander cells (**Figure 1G**). Some ISGs that are more highly upregulated by CL8-infected cells, including *MX2* and *IFITM1*, have known roles as HIV-1 restriction factors in cell lines^48–52^.

Several of the main immune mediators that are expressed within co-regulated gene modules overlap with those known to be important in the acute phase of SIV and HIV infection, including CCL2, TNF, and IL1B^53–55^. However, we also identify unique chemokine signatures that distinguish responses to lentiviral pathogenic variants. For example, we observe expression of neutrophil chemoattractants *CXCL1* and *CXCL2* primarily by CL8-infected animals. We hypothesize that these chemokines, which are not expressed by 170-infected animals, may play important roles in downstream immune signaling cascades; unfortunately, we are unable to profile such cascades in this dataset as we did not profile neutrophils. Additionally, we found that 170-infected animals were the predominant producers of *CXCL10* and *CXCL16*, which both operated in dense longitudinal networks of pro-inflammatory cytokine regulation. These findings implicate *CXCL10* and *CXCL16* in immune responses to highly pathogenic lentiviral infection and provide an avenue for future study.

Additionally, we used novel longitudinal frameworks for single-cell resolution CCC analysis to reveal that broad, sustained, and densely-connected regulatory networks of pro-inflammatory cytokines are a defining feature of responses to highly pathogenic lentiviral infection. While longitudinal cytokine regulatory networks were largely disconnected in both CL8-infected animals, in 170-infected animals pro-inflammatory cytokines were predicted to upregulate additional cytokines that were active at future time points. This is suggestive of amplifying positive feedback loops that result in the observed breadth of inflammatory activity in highly pathogenic lentivirus infection. While the distinct structure of these regulatory networks may be due to unique immune phenotypes induced by 170-infected cells, it is also possible that this observation is driven primarily by sustained viral loads in these animals.

Collectively, we perform the first single-cell resolution dissection of the distal immune mechanisms that distinguish infection with highly related lentiviral variants of different virulence. We show that cells infected with highly pathogenic lentivirus fail to upregulate the same breadth and magnitude of ISGs than cells infected with a less virulent virus as a potential immune escape mechanism and demonstrate that delayed and broad inflammatory pathways are characteristics of pathogenic lentiviral infection. We nominate potential drivers of these inflammatory pathways by using a combination of gene module discovery and novel longitudinal CCC analysis frameworks and reveal distinct regulatory programs of pro-inflammatory chemokines delineating responses to highly pathogenic lentiviral infection. Together, our work provides several mechanistic targets for further investigation that may represent targets for modulating lentiviral pathogenicity.

## ACKNOWLEDGEMENTS

We thank all current and former members of the Blish laboratory for helpful discussions of this work. A.J. Wilk is supported by the Stanford Medical Scientist Training Program (T32 GM007365-44) and the Stanford Bio-X Interdisciplinary Graduate Fellowship. This work was also supported by NIH/NIDA DP1 DA04508902 to C.A.B., NIH HD103571 to JO and CAB, NIH/NCI 1U54CA217377, U01 28020510, and 1U2CCA23319501 to A.K.S., NIH/NIDA 1DP1DA053731 to A.K.S., Bill and Melinda Gates Foundation INV-027498 and OPP1202327 to A.K.S., the MIT Stem Cell Initiative through Foundation MIT to A.K.S. and a 2019 Sentinel Pilot Project from the Bill and Melinda Gates Foundation to C.A.B./A.K.S..

## AUTHOR CONTRIBUTIONS

A.J.W., A.K.S., S.H., J.O., and C.A.B. conceived of the work and designed experiments. A.J.W., J.O.M., S.W.K., I.F., V.M., and J.G-R. performed scRNA-seq experiments. J.O.M. performed bulk RNA sequencing work. A.J.W. performed computational analyses. A.J.W. wrote the manuscript with input from all authors.

## DATA AVAILABILITY

Raw and processed scRNA-seq data generated in this manuscript will be deposited on GEO without restrictions.

## CODE AVAILABILITY

All original code used for analysis and visualization will be available from a Github repository.

## MATERIALS & METHODS

### Sample collection and storage

All PBMC samples profiled in this study were processed and collected as a part of a study by Kimata, et al.^12^. Briefly, juvenile pig-tailed macaques (*Macaca nemestrina*) were inoculated intravenously with equal amounts of SIVMneCL8 or SIVMne170. Serial samples of peripheral blood, serum, and plasma were collected prior to and weekly after inoculation to assess animal health, CD4^+^ T cell count, and viral load. Viral load measurements reported in this study were determined by Kimata, et al.^12^ by Chiron branch DNA (bDNA) assay. PBMCs were isolated by density centrifugation and cryopreserved. Samples from two animals infected with SIVMneCL8 and two animals infected with SIVMne170 were analyzed. From each animal, a pre-infection sample along with eight post-infection samples (from weeks 1, 2, 3, 4, 6, 8, 12, and 20 postinfection) were analyzed.

### Sample thawing and processing for transcriptomic analysis

PBMCs were thawed at 37°C in complete RPMI-1640 media (supplemented with 10% FBS, L-glutamine, and Penicillin-Streptomycin-Amphotericin; RP10) containing benzonase (EMD Millipore). Cells were then counted and diluted in RP10 to a concentration of 75,000 live cells/mL. Cell viability was determined by Trypan Blue staining and is reported in **Supplementary Figure 1**. Unless otherwise noted, all centrifugations were performed at 300 *x* g for 3 minutes. We observed that samples from both 170-infected animals at 1 week post-infection had extremely low viability post-thaw (18% and 8%, respectively). To attempt to rescue cell viability in these we applied two strategies for live cell enrichment. First, we performed soft centrifugation on these samples, at 100 *x* g for 1 minute, and discarded the supernatant. Next we applied the Dead Cell Removal Kit (catalog no. 130-090-101; Miltenyi BioTec) to the remaining cell pellet. This process improved viability to 88% and 80%, respectively, and we proceeded with downstream transcriptomic profiling.

### scRNA-seq by Seq-Well

The Seq-Well platform for scRNA-seq was utilized as described previously^13,22,56^. 200 μL of the cell suspension of thawed PBMCs at 75,000 live cells/mL (15,000 cells) was loaded onto Seq-Well arrays pre-loaded with mRNA capture beads (ChemGenes). Following four washes with serum-free RPMI-1640 to remove serum, the arrays were sealed with a polycarbonate membrane (pore size of 0.01 μm) for 30 minutes at 37°C. Next, arrays were placed in lysis buffer, transcripts hybridized to the mRNA capture beads, and beads recovered from the arrays and pooled for downstream processing. Immediately after bead recovery, mRNA transcripts were reverse transcribed using Maxima H-RT (Thermo Fisher EPO0753) in a templateswitching-based RACE reaction, excess unhybridized bead-conjugated oligonucleotides removed with Exonuclease I (NEB M0293L), and second-strand synthesis performed with Klenow fragment (NEB M0212L) to enhance transcript recovery in the event of failed template switching ^14^. Whole transcriptome amplification (WTA) was performed with KAPA HiFi PCR Mastermix (Kapa Biosystems KK2602) using approximately 6,000 beads per 50 μL reaction volume. Resulting libraries were then pooled in sets of 6 (approximately 36,000 beads per pool) and products purified by Agencourt AMPure XP beads (Beckman Coulter, A63881) with a 0.6x volume wash followed by a 0.8x volume wash. Quality and concentration of WTA products was determined using an Agilent Fragment Analyzer (Stanford Functional Genomics Facility), with a mean product size of >800bp and a non-existent primer peak indicating successful preparation. Library preparation was performed with a Nextera XT DNA library preparation kit (Illumina FC-131-1096) with 1 ng of pooled library using dual-index primers. Tagmented and amplified libraries were again purified by Agencourt AMPure XP beads with a 0.6x volume wash followed by a 1.0x volume wash, and quality and concentration determined by Fragment Analysis. Libraries between 400-1000bp with no primer peaks were considered successful and pooled for sequencing. Sequencing was performed on a NovaSeq 6000 instrument (Illumina; Chan Zuckerberg Biohub). The read structure was paired-end with read 1 beginning from a custom read 1 primer^13^ containing a 12bp cell barcode and an 8 bp unique molecular identifier (UMI), and with read 2 containing 50bp of mRNA sequence.

### Bulk RNA-sequencing

PBMCs were removed from liquid nitrogen and thawed on ice. A subset of cells were counted and removed for Seq-Well analysis, while the remaining cells were pelleted and immediately disrupted with Buffer RLT (Qiagen). Extraction and purification of total RNA from bulk PBMCs was conducted using Qiagen RNeasy plus kit (#74034). All extractions included genomic DNA removal using Qiagen gDNA Eliminator Spin Columns. The concentration of RNA was quantified using a Nanodrop 2000 Spectrophotometer (ThermoFisher). RNA quality was assessed using RIN values obtained from Agilent 4200 Tapestation analysis. Reverse transcription and tagmentation were conducted using the SMART-Seq v4 (Takara) and Nextera XT library prep kit (Illumina) with an input of 10ng of RNA per sample. After cDNA synthesis and library preparation, the samples were sequenced using Illumina NovaSeq SP with paired-end 50-bp read length, yielding approximately 30M reads per sample.

Alignment and quality control of single-cell RNA sequencing data Sequencing reads were aligned and count matrices assembled using STAR^57^ and dropEst^58^, respectively. Briefly, the mRNA reads in read 2 demultiplexed FASTQ files were tagged with the cell barcode and UMI for the corresponding read in the read 1 FASTQ file using the dropTag function of dropEst. Next, reads were aligned with STAR using the *M. nemestrina* reference genome Mnem_1.0 that included the sequences of the SIVMneCL8 and SIVMne170 genomes for alignment. Given the similarity between the SIVMneCL8 and SIVMne170 genomes, any read aligning to either SIV genome sequence was annotated as an SIV viral read. Any cell with >0 UMIs aligned to the SIV genome is considered SIV^+^. Count matrices were built from resulting BAM files using dropEst^58^. We noted that several Ensembl gene IDs had annotations available that were not present in the GTF from the latest Mnem_1.0 release (**Supplementary Figure 1**). We transferred these additional annotations to the count matrices using the R package biomaRt^59^. Cells that had fewer than 1,000 UMIs or greater than 15,000 UMIs, as well as cells that contained greater than 20% of reads from rRNA genes (*RNA18S5* or *RNA28S5*) or greater than 0.1% of reads from one of the 6 annotated mitochondrial genes in the *M. nemestrina* genome, were considered low quality and removed from further analysis. To remove putative multiplets (where more than one cell may have loaded into a given well on an array), cells that expressed more than 75 genes per 100 UMIs were also filtered out. Genes that were expressed in fewer than 10 cells were removed from the final count matrix.

### Pre-processing of single-cell RNA sequencing data

The R package Seurat^60–62^ was used for data scaling, transformation, clustering, dimensionality reduction, differential expression analysis, and most visualizations. Data were scaled and transformed and variable genes identified using the SCTransform() function, and linear regression performed to remove unwanted variation due to cell quality (% mitochondrial reads, % rRNA reads). PCA was performed using the 3,000 most highly variable genes. The first 50 principal components (PCs) were used to perform dimensionality reduction by UMAP^63,64^, to construct a shared nearest neighbor graph (SNN; FindNeighbors()), and this SNN used to cluster the dataset (FindClusters()).

### Automated annotation of granular cell types by Seurat v4

We used the multimodal (whole transcriptome plus 228 cell surface proteins) PBMC dataset published by Hao, et al.^62^ as a reference for cell type annotation. We first scaled both the transcriptome and protein assays, and ran PCA on both modalities. Next, we found multimodal neighbors between the modalities via weighted nearest neighbor (WNN) analysis, which learns the relative utility of each data modality in each cell. Supervised PCA (SPCA) was then run on the WNN SNN graph, which seeks to capture a linear transformation that maximizes its dependency to the WNN SNN graph. These SPCA reduced dimensions were then used for identification of anchors between the reference and query datasets as previously described^60^. Finally, these anchors were used to transfer reference cell type annotations to the query dataset. As the Hao, et al. reference dataset used for annotation is a human dataset, we confirmed the validity of the transferred annotations by calculating differentially expressed genes for each cell type annotation using Seurat’s implementation of the Wilcoxon rank-sum test (FindMarkers()) and comparing those markers to known cell type-specific genes from previous datasets^65–70^. Cell type annotations transferred with this strategy matched biological expectations and overlapped with results from graph-based clustering (**Supplementary Figure 1**).

### Calculation of transcriptomic perturbation score

To prioritize analysis of cell types of interest, we calculated a perturbation score for each cell type of each sample as previously described^71,72^. This perturbation score is motivated by the observation that the statistical significance of per-gene differential expression tests is strongly influenced by the number of cells in each cluster or cell type. To overcome this, we first identified a set of genes for each cell type that showed evidence of differential expression (p value < 0.1) between a post-infection and pre-infection time point. Next, we defined the global perturbation vector as the average log-fold change of each DEG relative to pre-infection samples normalized to length 1. Finally, we projected the transcriptome of each sample onto this vector and defined the perturbation score as the absolute value of the magnitude of this projection. This approach enables prioritization of cell type perturbations when comparing cell types of different abundances.

### Gene module scoring analysis

The Seurat function AddModuleScore() was used to score single cells by expression of a list of genes of interest. This function calculates a module score by comparing the expression level of an individual query gene to other randomly-selected control genes expressed at similar levels to the query genes, and is therefore robust to scoring modules containing both lowly and highly expressed genes, as well as to scoring cells with different sequencing depth. Gene lists for module scoring were generated through gene module discovery. A previously published list of ISGs was used to define the interferon signature score (**Supplementary Table 2**)^73^.

### Differential expression testing

We used Seurat’s implementation of differential expression testing in FindMarkers() to identify DE genes and DE ligand-receptor pairs. Unless otherwise specified, testing for DE genes was performed via Seurat’s implementation of the Wilcoxon rank-sum test, which reports two-sided p-values corrected for multiple hypothesis testing by Bonferroni’s correction. For single-cell resolution CCC analysis with Scriabin, ligand-receptor pair activity values are not independent observations and therefore a receiver operating characteristic curve (ROC) test was used for differential expression testing. In this test, a classifier is built on each gene which is used to classify two groups of cells being compared. An AUC value of 1 indicates that the expression values for that gene can perfectly classify the two cell groups.

Differential expression testing of ISGs (**Figure 2C-D**) was performed on a per animal per cell type basis: for each sample, DE genes were calculated between each post-infection time point relative to the respective pre-infection time point. Cumulative log(fold-changes) represent the sum of the fold-changes for each post-infection time point at which the gene was significantly (p-value < 0.05) upregulated. For **Figures 2E-F**, only genes that were significantly DE (p-value < 0.05) in at least one of the analyzed dimensions were plotted for visualization.

### Gene module discovery with WGCNA

Unless otherwise noted, gene module analysis was performed as previously described^22^. This process was performed separately for each major cell type from each animal. Briefly, principal components (PCs) of biological interest were first identified by examining the variance structure described by each PC and the genes contributing to each PC. PCs used for each WGCNA analysis are shown in **Supplementary Table 3**. Variable genes of biological interest were defined as the top and bottom 50 genes for each PC and the input to WGCNA was subset to these genes.

Next, a signed genes covariance matrix is calculated for each sample:

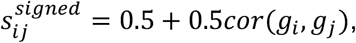

where *g_i_*,*g_j_* are individual genes. 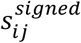 is next converted into an adjacency matrix via soft thresholding:

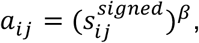

where *β* is the soft power. Soft power is a user-defined parameter that is recommended to be the lowest value that results in a scale-free topology model fit of > 0.8^23^. Soft power parameters used for analysis are shown in **Supplementary Table 3**. Next, this adjacency matrix is converted into a TOM as described^74^. Next, this TOM was hierarchically clustered and cutreeDynamic used to generate modules of highly correlated genes with a minimum module size of 10. Similar modules were merged using a dissimilarity threshold of 0.5. Subsequently, the correlation structure of each module was tested using a permutation test, comparing the correlation of randomly generated modules to the true module. A module was considered significantly correlated if < 500 of 10,000 one-sided Mann-Whitney *U*-tests were significant at the p < 0.05 level.

Single cells were then scored for expression of each module as described above. Finally, we tested for modules with significant variation in expression over time. Since testing for differences in distribution is highly sensitive to sample size, we performed iterations of a Kruskal-Wallis test for expression of each module relative to time point on 10,000 equally-sized cell subsets. Modules were considered significantly temporally-variable if < 500 of 10,000 Kruskal-Wallis tests were significant at the p < 0.05 level.

### Analysis of CCC with Scriabin

Scriabin^30^ was used for analysis of CCC at single-cell resolution as previously described. In brief, the center of Scriabin’s workflow is the generation of a cell-cell interaction matrix (CCIM) that captures the full structure of CCC phenotypes in the dataset. For a dataset of *n* cells and *k* possible ligand-receptor pairs that can be used for CCC, the CCIM is a matrix of *n* × *n* columns of single cell-cell pairs by *k* ligand-receptor pairs where each matrix element represents the predicted activity of a given ligand-receptor pair between a cell-cell pair.

#### Generation of summarized interaction graphs

Because the CCIM scales exponentially with dataset size, it was impractical to calculate a CCIM for all cell-cell pairs *n_i_*,*n_j_*. To address this problem, a summarized cell-cell interaction graph ***S*** was built in lieu of the CCIM where an element of ***S*** is the sum of all potential ligand-receptor activities between a cell-cell pair. A matrix ***S*** was built for each time point.

#### Dataset binning for comparative CCC analyses

Conceptually, comparing ***S*** from multiple samples requires that cells from different samples share a set of labels or annotations denoting what cells represent the same identity. Each identity class to be compared then requires representation from each of the samples to be compared. This inter-dataset alignment was accomplished through Scriabin’s binning workflow. The goal of binning is to assign each cell a bin identity so that ***S*** from multiple samples can be summarized into equidimensional matrices based on shared bin identities.

First, a shared nearest neighbor (SNN) graph was constructed via FindNeighbors() to define connectivity between all cells to be compared. Next, mutual nearest neighbors (MNNs) were identified between all time points to be compared via Seurat’s integration workflow (FindIntegrationAnchors())^60^. Next, bin identities were assigned based on these inter-dataset MNN pairings and iteratively optimized for the SNN connectivity of cells within the same bin and for the number of time points represented within each bin. Finally, incompletely represented bins were merged together based on SNN connectivity. At the end of this process, each cell has a single assigned bin identity, where each bin contains cells from all samples to be compared.

Bins were then tested for statistical significance of connectivity structure using a permutation test. For each bin, random bins of the same size and number of cells per sample were generated iteratively (by default 10,000 times). The connectivity vector of the real bins was tested against each of the random bins by a one-sided Mann-Whitney U test. If the bin failed 500 or more of these tests (p-value 0.05), it was considered non-significant, discarded, and merged with its most similar counterpart.

Finally, bin-bin pairs that were variable in their CCC magnitude over time were identified. For each bin-bin pair, a Kruskal-Wallis test was used to assess differences in the magnitude of CCC between cell-cell pairs from different time points. Cells from bin-bin pairs with a two-sided Kruskal-Wallis p-value < 0.05 and within the top 75 percentile of Kruskal-Wallis test statistic were considered highly perturbed. Dunn’s post-hoc test was used to identify the most frequently perturbed time point among highly variable bin-bin pairs. Cells from highly variable bins in cell type combinations of interest were then used to construct CCIMs.

#### Identification of biologically-active ligand-receptor edges

To identify what ligands were predicted to be active in each receiver cell, we first calculated a per-cell gene signature of temporally-variable genes using the package CelliD^75^. Next, potential ligands were defined as those ligands that were expressed by at least 2.5% of sender cells being analyzed. The ligand activity for each receiver cell was defined as the Pearson correlation coefficient between each cell’s gene signature, and the vector of ligand-to-target gene links for that ligand^76^. For each active ligand, target gene weights for each cell were defined as the ligand-target matrix regulatory score for the top 250 targets for each ligand that appear in a given cell’s gene signature. We selected a Pearson coefficient threshold (by default 0.075) to define active ligands in each cell.

To identify statistically significant changes in ligand activity over time, we performed iterations of a Kruskal-Wallis test for expression of each ligand activity score relative to time point on 10,000 equally-sized cell subsets. Ligand activities were considered significantly temporally-variable if < 500 of 10,000 Kruskal-Wallis tests were significant at the p < 0.05 level.

#### CCIM construction

We defined the interaction vector *V* between sender cell *N_i_* and receiver cell *N_j_* as:

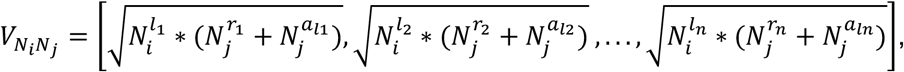

where *l_n_*,*r_n_* represent a cognate ligand-receptor pair and *α_ln_* represents the predicted activity of an active ligand *l_n_* in *N_j_*. The CCIM was assembled by concatenating interaction vectors *V_N_i___N_j__* for all cell-cell pairs. Individual CCIMs were generated for each sample, so that cell-cell pairs were only generated between cells that could be biologically interacting.

#### Downstream analysis of CCIM

The CCIM can be treated analogously to the gene expression matrix and used for downstream analysis tasks like dimensionality reduction. To focus our downstream analysis on ligand-receptor pairs of biological interest, we subset the CCIM to ligand-receptor pairs where either the ligand or receptor appeared within the top 1,000 genes of the parent object. To remove additional sources of noise in downstream visualization, we also discarded cell-cell pairs where fewer than 5 ligand-receptor pairs were non-zero in activity. Next, we scaled this object by ScaleData(), found latent axes by RunPCA(), and used the top 15 PCs to embed the dataset in two dimensions using UMAP^64^. Neighbor graphs were constructed by FindNeighbors(), which were then clustered via modularity optimization graph-based clustering^77^ as implemented by Seurat’s FindClusters()^62^.

### Identification of longitudinal CCC circuits

Longitudinal communication circuits were identified as previously described^30^. Briefly, a longitudinal CCC circuit is composed of S_1_-L_1_-R_1_-S_2_-L_2_-R_2_, where S are sender cells and R are receiver cells at timepoints 1 and 2, and where L_1_ is expressed by/sensed by S_1_/R_1_ and L_2_ is expressed by/sensed by S_2_/R_2_. Ligand activities and target gene linkages used for circuit generation were calculated as described above.

### Visualization

Wrappers provided by Seurat were used to generate UMAP projection and Dot Plots. ComplexHeatmap was used to generate all heatmaps. Visualization functions available from Scriabin were used to visualize most cell-cell communication pathways. Custom ggplot functions (see **Data and materials availability**) were used to generate all other plots. For all boxplots, features include: minimum whisker, 25th percentile–1.5 × inter-quartile range (IQR) or the lowest value within; minimum box, 25th percentile; center, median; maximum box, 75th percentile; maximum whisker, 75th percentile + 1.5 × IQR or greatest value within. Feature line plots are boxplots where consecutive time points from the same animal are connected via a straight line. Unless otherwise noted, all feature line plots contain a 95% confidence interval around each point.

