## Supplementary Information for "Pro-inflammatory feedback loops define immune responses to pathogenic lentivirus infection"

**This PDF file includes:**

Supplementary Tables 1-4
Supplementary Figures 1-8

**Supplementary Tables**

| **ENSEMBL ID** | **Renamed ID** |
|---|---|
| ENSMNEG00000030800 | CCL3L |
| ENSMNEG00000034156 | CCL3 |
| ENSMNEG00000040223 | CCL4 |
| ENSMNEG00000031998* | IFITM3 |
| ENSMNEG00000038291* | IFI27-2 |
| ENSMNEG00000027314 | FAM127A |
| ENSMNEG00000039432 | HLA-C |
| ENSMNEG00000043296 | MANE-B-2 |
| ENSMNEG00000033920 | HLA-DQA1-2 |
| ENSMNEG00000044119 | HLA-DRB1 |
| ENSMNEG00000044507 | HLA-DMB |
| ENSMNEG00000039329 | FCGR3 |

**Supplementary Table 1: List of manually-annotated ENSEMBL IDs.** Asterisks indicate ENSEMBL IDs that were included for IFN signature scoring and analysis.

| Interferon-stimulated gene | | | | | |
|---|---|---|---|---|---|
| ADAR | DDX60 | IFITM1 | MVB12A | PSMB8 | TENT5A |
| B2M | DHX58 | IL15 | MX1 | PSMB9 | TRAFD1 |
| BATF2 | EIF2AK2 | IL4R | NCOA7 | PSME1 | TRIM14 |
| BST2 | ELF1 | IL7 | NMI | PSME2 | TRIM21 |
| C1S | EPSTI1 | IRF1 | NUB1 | RIPK2 | TRIM25 |
| CASP1 | GMPR | IRF2 | OAS1 | RNF31 | TRIM26 |
| CASP8 | HELZ2 | IRF7 | OASL | RSAD2 | TXNIP |
| CCRL2 | HERC6 | IRF9 | OGFR | RTP4 | UBA7 |
| CD47 | IFI27 | ISG15 | PARP12 | SAMD9 | UBE2L6 |
| CD74 | IFI30 | ISG20 | PARP14 | SELL | IFI6 |
| CMPK2 | IFI35 | LAMP3 | PARP9 | SLC25A28 | MX2 |
| CMTR1 | IFI44 | LAP3 | PLSCR1 | SP110 | GBP3 |
| CNP | IFI44L | LGALS3BP | PNPT1 | STAT2 | OAS2 |
| CSF1 | IFIH1 | LPAR6 | PROCR | TAP1 | IFIT5 |
| CXCL10 | IFIT2 | LY6E | PSMA3 | TDRD7 | GBP7 |
| CXCL11 | IFIT3 | MOV10 | IFI16 | | |

**Supplementary Table 2: List of ISGs.**

| Cell type | nPC | SoftPower | Animal |
|---|---|---|---|
| B | 30 | 6 | 170 A |
| B | 30 | 4 | 170 B |
| B | 30 | 3 | CL8 A |
| B | 30 | 6 | CL8 B |
| CD14 Mono | 29 | 7 | 170 A |
| CD14 Mono | 29 | 5 | 170 B |
| CD14 Mono | 30 | 6 | CL8 A |
| CD14 Mono | 21 | 6 | CL8 B |
| CD16 Mono | 30 | 6 | 170 A |
| CD16 Mono | 20 | 5 | 170 B |
| CD16 Mono | 30 | 5 | CL8 A |
| CD16 Mono | 30 | 5 | CL8 B |
| CD4 T | 30 | 5 | 170 A |
| CD4 T | 30 | 7 | 170 B |
| CD4 T | 30 | 6 | CL8 A |
| CD4 T | 30 | 6 | CL8 B |
| CD8 T | 30 | 3 | 170 A |
| CD8 T | 29 | 3 | 170 B |
| CD8 T | 30 | 3 | CL8 A |
| CD8 T | 30 | 5 | CL8 B |

| | | | |
|---|---|---|---|
| DC | 25 | 3 | 170 A |
| DC | 25 | 6 | 170 B |
| DC | 25 | 5 | CL8 A |
| DC | 25 | 7 | CL8 B |
| NK | 30 | 5 | 170 A |
| NK | 28 | 4 | 170 B |
| NK | 29 | 5 | CL8 A |
| NK | 30 | 5 | CL8 B |

**Supplementary Table 3: Parameters used for WGCNA-based gene module discovery.**

| Pro-inflammatory cytokine gene | | | |
|---|---|---|---|
| IL18 | TNF | CCL15 | CCL8 |
| IL18BP | TNFSF10 | CCL16 | CX3CL1 |
| IL1A | TNFSF11 | CCL17 | CXCL1 |
| IL1B | TNFSF12 | CCL18 | CXCL10 |
| IL36A | TNFSF13 | CCL19 | CXCL11 |
| IL37 | TNFSF15 | CCL2 | CXCL12 |
| IL1RL2 | TNFSF4 | CCL20 | CXCL13 |
| IL33 | IFNB1 | CCL22 | CXCL14 |
| TNFSF9 | IFNG | CCL25 | CXCL16 |
| TNFSF8 | CLCF1 | CCL28 | CXCL17 |
| FASLG | IL31 | CCL3 | CXCL2 |
| TNFSF14 | IL6 | CCL4 | CXCL6 |
| LTA | LEPR | CCL5 | CXCL9 |
| LTB | LIF | CCL7 | CXCL8 |
| | | | XCL1 |

**Supplementary Table 4: List of pro-inflammatory cytokine-encoding genes.**

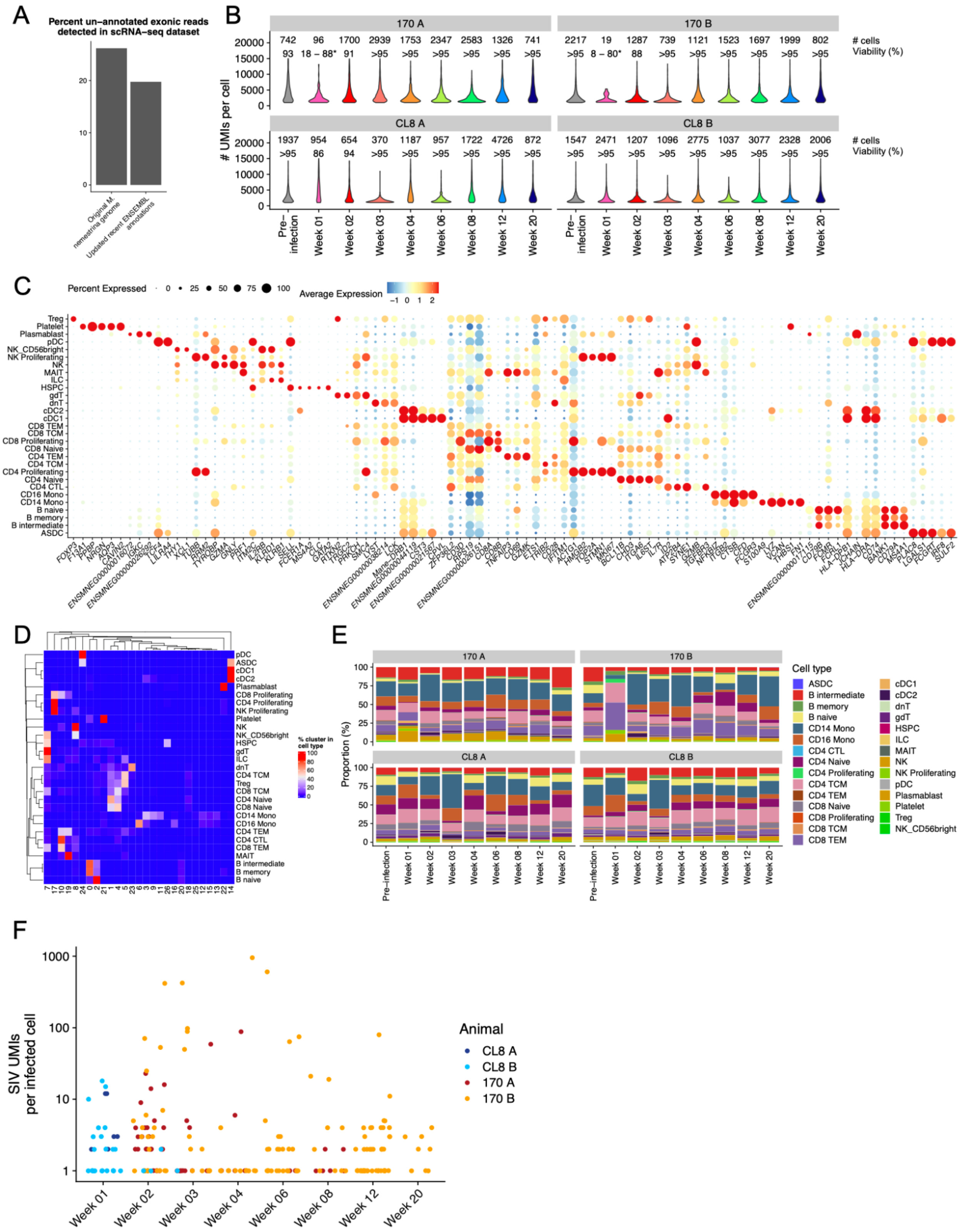

**Supplementary FIgure 1: Quality control of single-cell transcriptomic dataset**. **A)** Bar plot depicting the percentage of un-annotated exonic entries in the original Mnem_1.0 reference genome and after updating with recent annotation additions from Ensembl. **B)** Violin plots depicting the number of UMIs per cell per sample. The text above each violin shows the number of cells from each sample (top) and the viability of the sample after thawing (bottom). PBMCs from 170-infected animals at 1 week post-infection thawed with low viability (18% and 8%, respectively) and we applied dead cell removal techniques (see **Methods**) to attempt to rescue cell viability. While post-dead cell removal viability dramatically improved (to 88% and 80%, respectively), very few high quality cells were sequenced from these samples, indicative of active cell death. **C)** Dot plot depicting percent and average expression of the top five DE genes for each cell type label from reference-based annotations (see **Methods**). **D)** Heatmap depicting overlap between Seurat v4-based annotations shown in (**C**) and graph-based clustering results. Each row sums to 100%. **E)** Stacked bar plots depicting the cell type composition profile of each sample using reference-based cell type annotations. **F)** Scatter plot depicting the number of SIV UMIs in each SIV$^+$ cell across each time point. Points are colored by the animal of origin.

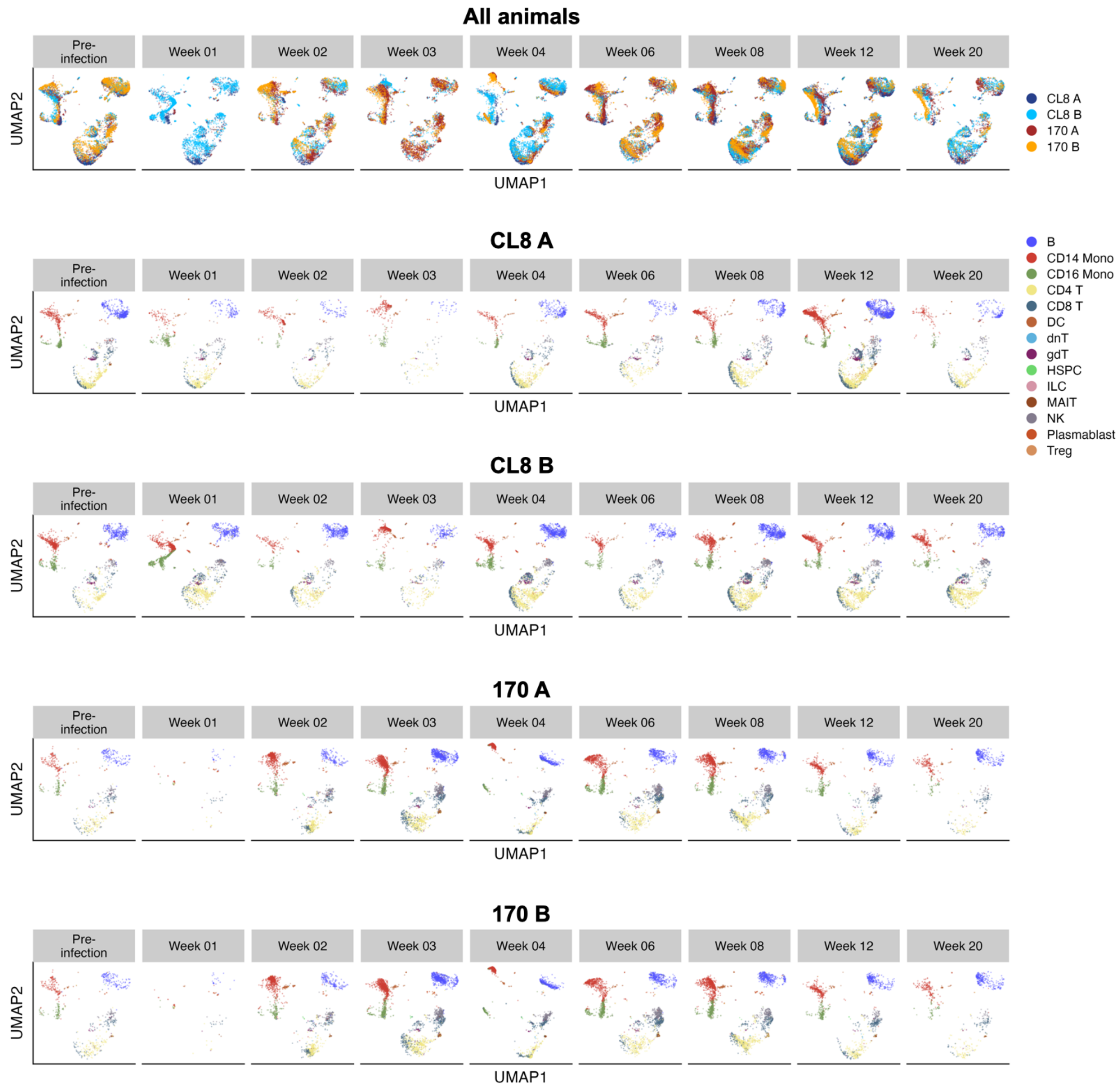


**Supplementary Figure 2: Dimensionality reduction embeddings faceted by time and animal**. The UMAP dimensionality reduction projection shown in **Figure 1D** is faceted by the time point from which each cell originated. In the top set of plots, cells from all animals are shown together and colored by the animal of origin. In the bottom four plots, the projections are also faceted by the animal of origin and colored by cell type.

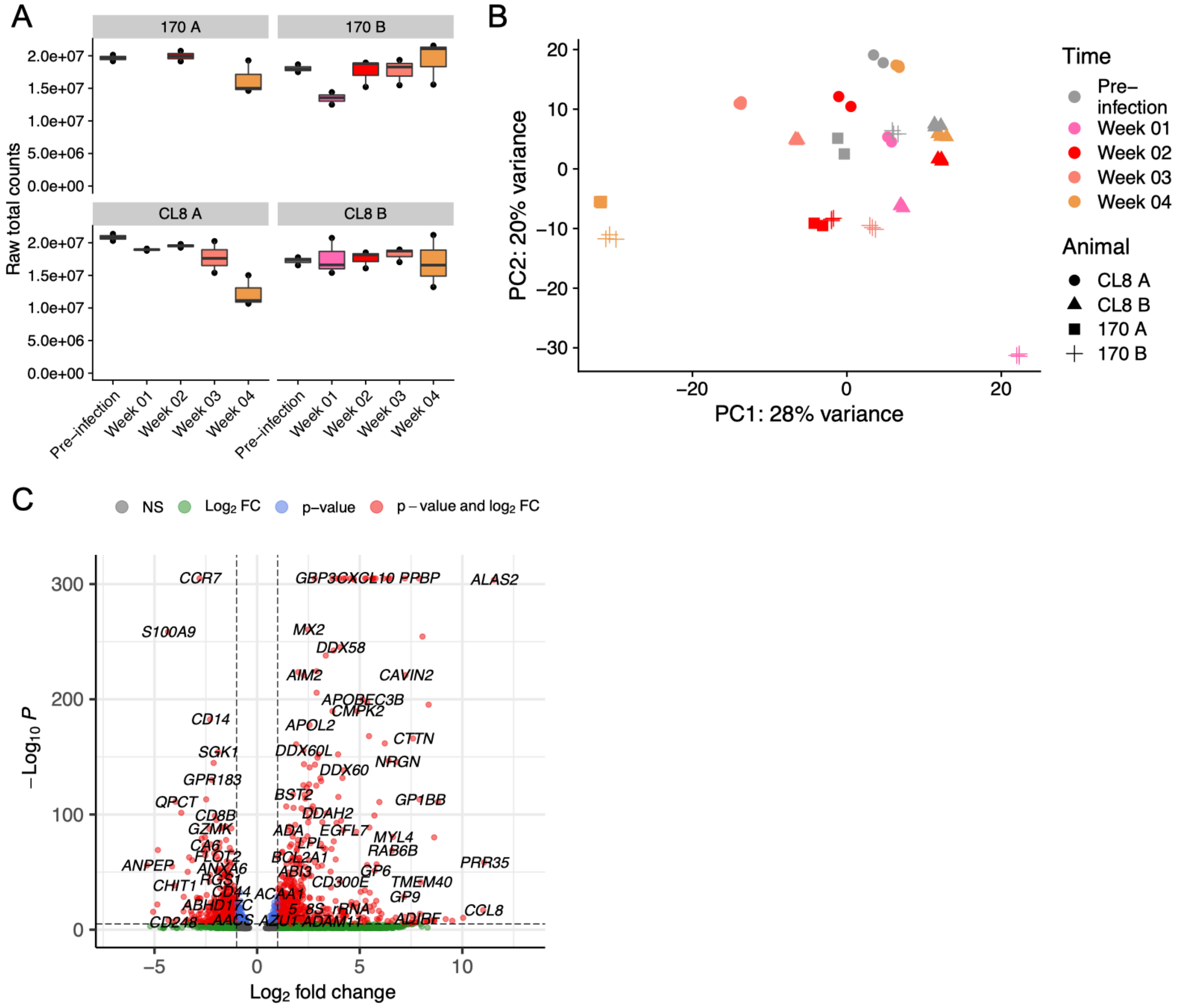


**Supplementary Figure 3: Comparison of bulk and single-cell transcriptomic profiles. A)** Box plots depicting total RNA counts from each sample profiled by bulk RNA sequencing. Each sample was processed either in duplicate or triplicate technical replicates. **B)** Projection of first two PCs of all samples profiled by bulk RNA sequencing. **C)** Volcano plot of DE genes in bulk RNA sequencing data, comparing animal 170 B 1 week post-infection (positive fold-change) to pre-infection (negative fold-change).

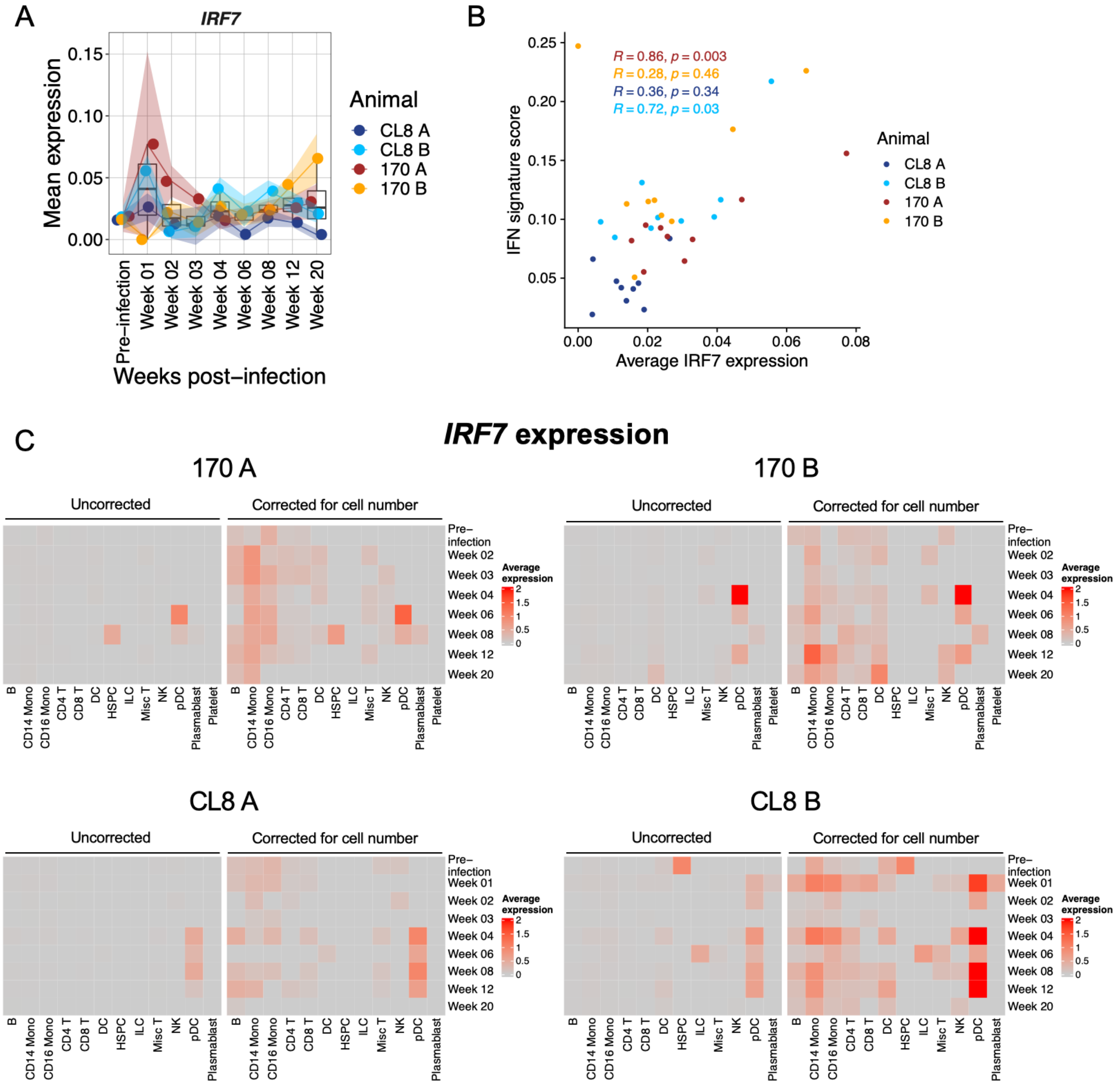


**Supplementary Figure 4: Expression of *IRF7* implicates sources of type I IFN. A)** Feature line plot (see **Methods**) depicting global expression of *IRF7* in all samples. **B)** Scatter plot depicting correlation between global *IRF7* expression and the IFN signature score. Pearson's r and exact two-sided p values are shown for the correlation for each animal. **C)** Heatmaps depicting the average expression of *IRF7* by each cell type at each time point. Left: raw average expression in each cell type. Right: average expression values were multiplied by the number of cells from the respective cell type at the respective time point.

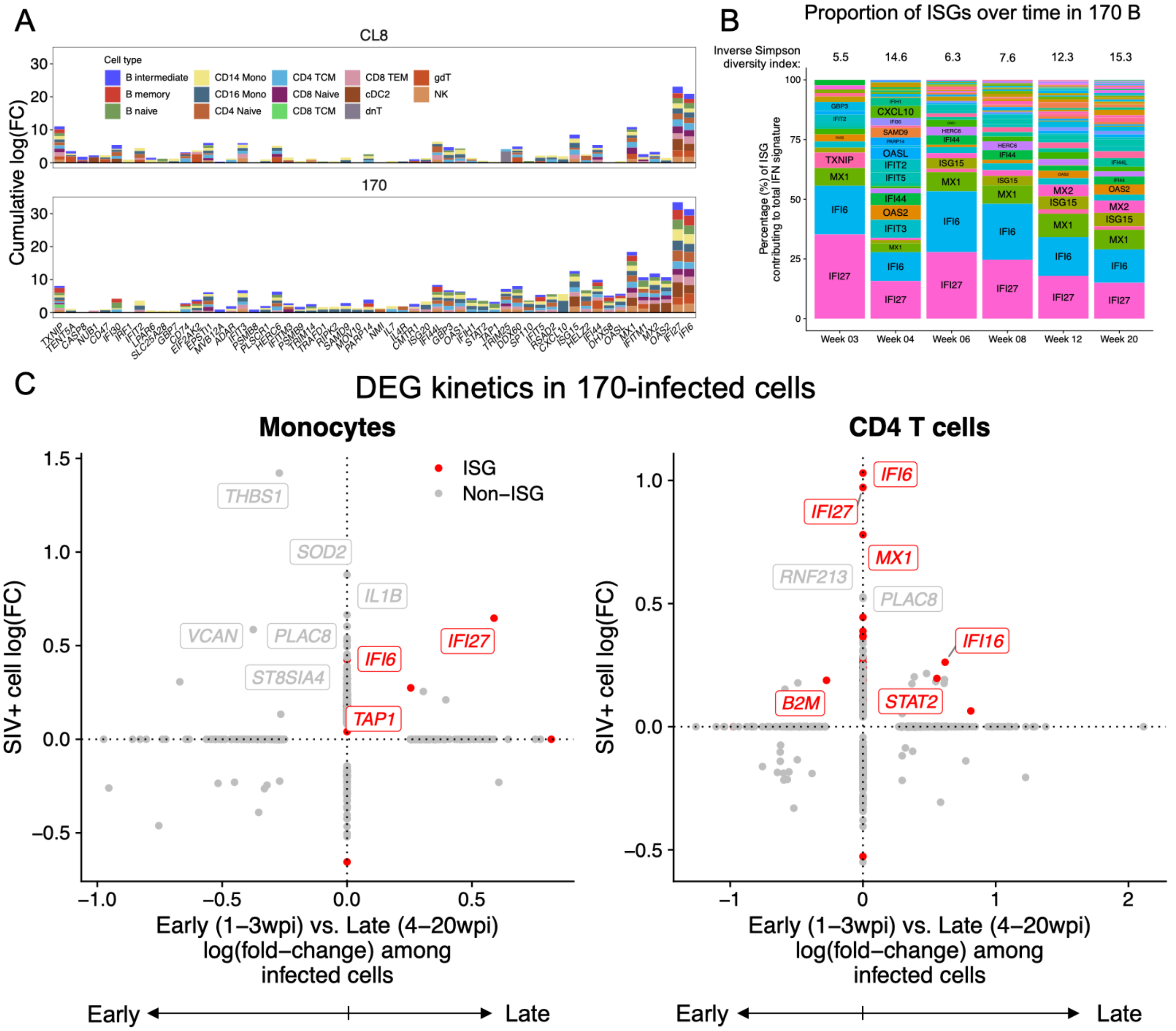


**Supplementary Figure 5: Analysis of IFN response kinetics. A-B)** For each animal, within each cell type, and at each non-baseline time point, differentially-expressed ISGs were calculated relative to pre-infection cells. **A)** For each ISG, 170-infected animal, and cell type, we calculated the cumulative log(fold-change) of significantly upregulated ISGs across all time points. **B)** For animal 170 B, we calculated the relative contribution of each ISG in the total IFN signature at each time point. The inverse Simpson index of diversity is plotted above each time point. **C)** In monocytes (left) and CD4$^{+}$ T cells (right), we calculated DEGs between SIV$^{+}$ cells relative to SIV$^{-}$ cells (*y*-axis). Then, among the 170-infected SIV$^{+}$ cells we calculated DEGs between cells from early time points (1-3 weeks post-infection; negative *x*-axis) vs. later time points (4-20 weeks post-infection; positive *x*-axis). All genes shown are significantly differentially expressed along at least one axis by Seurat's implementation of the Wilcoxon rank-sum test.

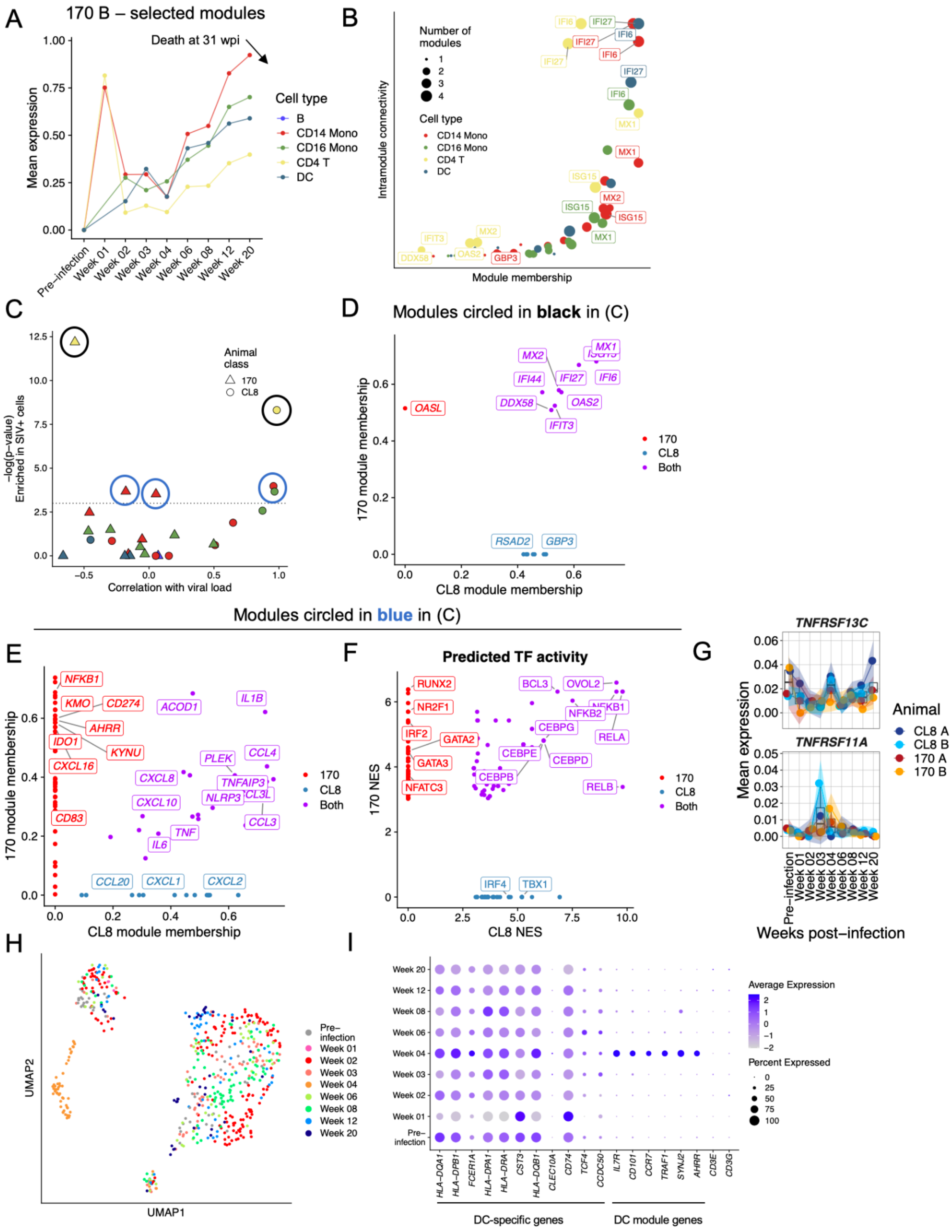


**Supplementary Figure 6: Analysis of gene modules correlated with viral load or enriched in SIV$^+$ cells. A)** Expression of gene modules in 170 B that monotonically increase in expression preceding death at 31 weeks post-infection. **B)** Scatter plot depicting module membership and intramodular connectivity of genes depicted in (**A**). The color of the points corresponds to the cell type in which the gene module was discovered and the size of the points corresponds to the number of modules in (**A**) to which each gene belongs. **C)** Scatter plot depicting module expression correlation with viral load (Pearson's r; *x*-axis) and enrichment of module expression in $SIV^+$ cells (-log(p-value) from Seurat's implementation of the Wilcoxon rank-sum test; *y*-axis). Modules are colored by the cell type in which the module was discovered and shapes correspond to the class of animal in which each module was discovered. **D)** Scatter plot depicting module membership for CL8- vs. 170-infected animals for the gene modules circled in black in (**C**). **E-F)** Scatter plots depicting module membership (**E**) and TF activity predictions (**F**) for CL8- vs. 170-infected animals for the gene modules circled in blue in (**C**). **G)** Feature line plots depicting global expression of receptors that induce non-canonical NF-κB signaling *TNFRSF13C* (encoding BAFFR) and *TNFRSF11A* (encoding RANK). **H)** UMAP projection of DCs from 170-infected animals colored by time point of sampling. **I)** Dot plot depicting percent and average expression of canonical DC marker genes, selected genes from the modules analyzed in **Figure 3G**, and canonical T cell marker genes.

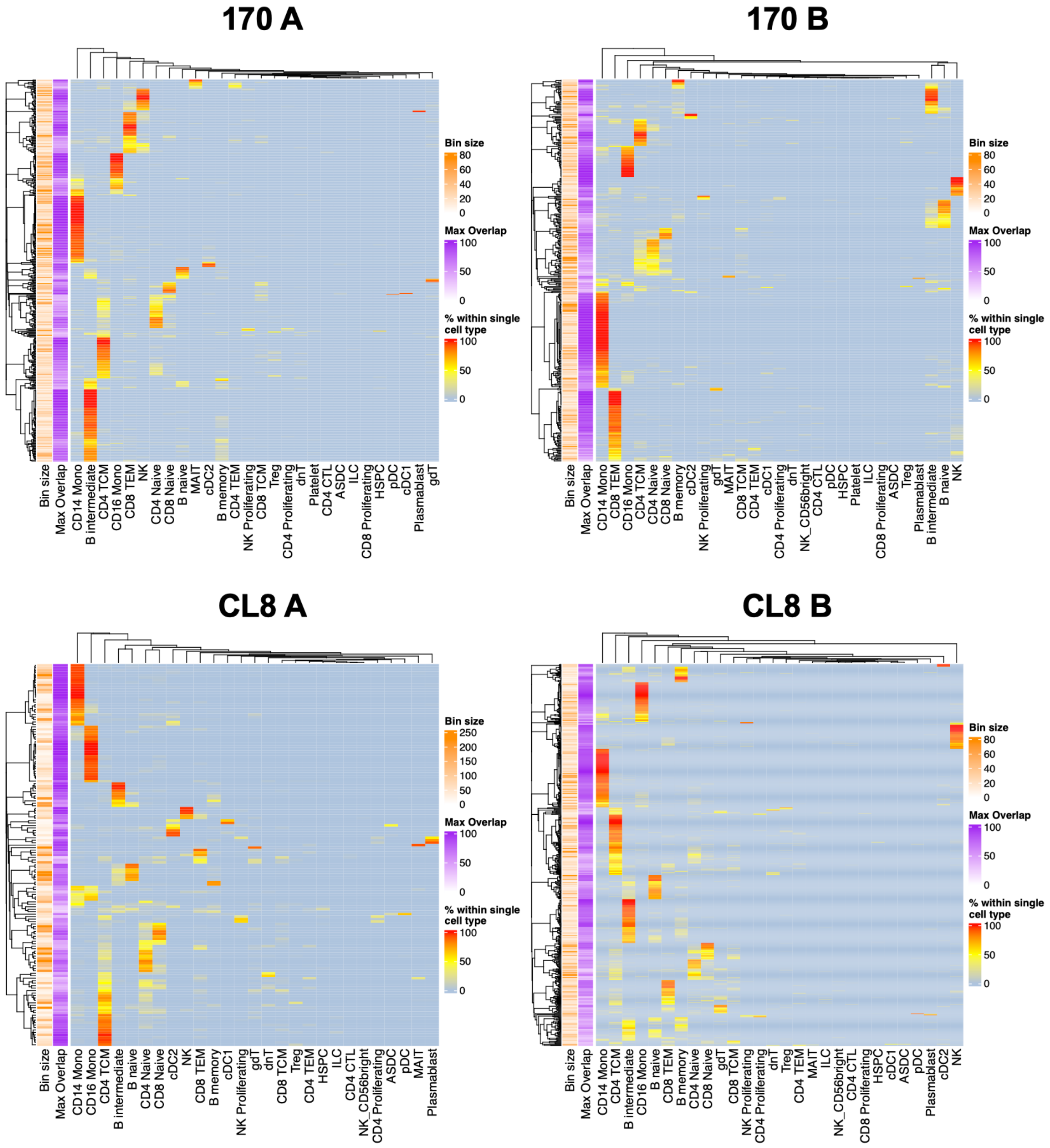


**Supplementary FIgure 7: Verification of binning fidelity.** Heatmaps of the overlap between cell bin identity and orthogonally-generated cell type annotations. Row annotations at the left of each heatmap correspond to the number of cells within each bin (left) and the percentage of cells belonging to the most represented cell type in each bin (the maximum value in each row; right). Each row sums to 100%.

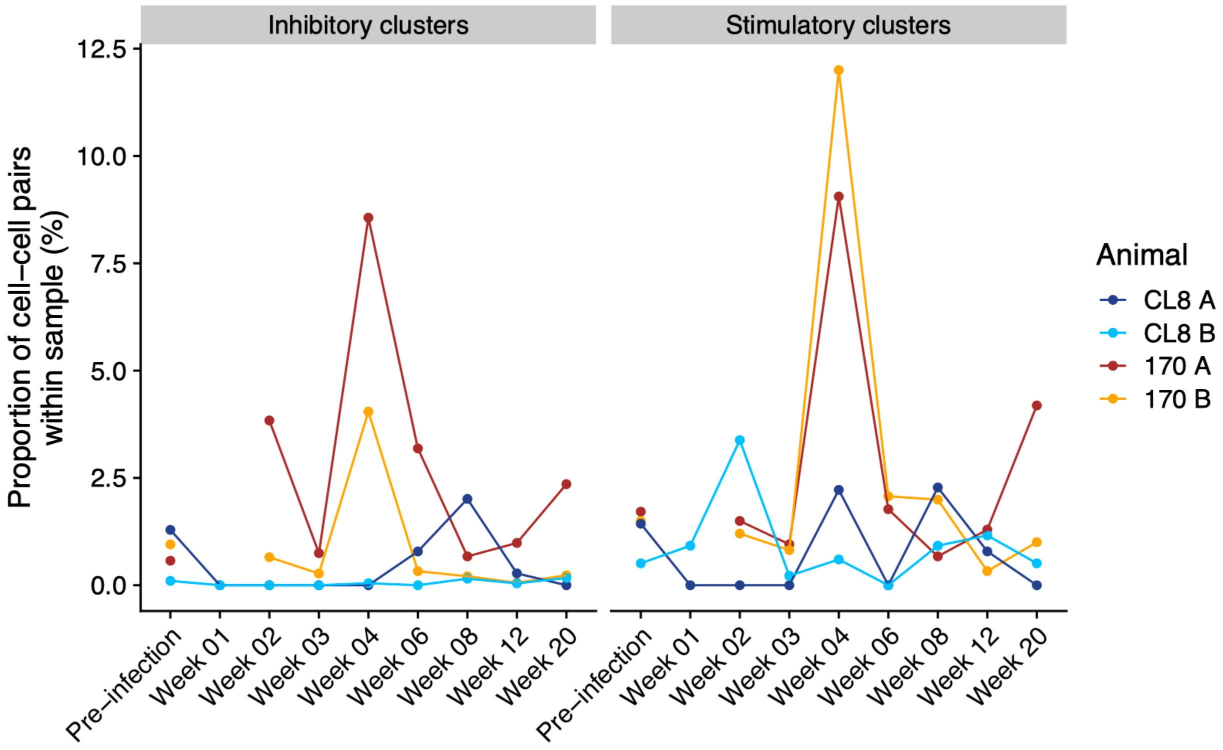


**Supplementary Figure 8**. **Abundance of inhibitory and stimulatory CCIM clusters in CD8 $T_{EM}$ to monocyte communication.** Proportion of cell-cell pairs from each sample that belong to the inhibitory (left) or stimulatory (right) clusters highlighted in **Figure 4H**.
